# Generalizing HMMs to Continuous Time for Fast Kinetics: Hidden Markov Jump Processes

**DOI:** 10.1101/2020.07.28.225052

**Authors:** Zeliha Kilic, Ioannis Sgouralis, Steve Pressé

## Abstract

The hidden Markov model (HMM) is a framework for time series analysis widely applied to single molecule experiments. It has traditionally been used to interpret signals generated by systems, such as single molecules, evolving in a discrete state space observed at discrete time levels dictated by the data acquisition rate. Within the HMM framework, originally developed for applications outside the Natural Sciences, such as speech recognition, transitions between states, such as molecular conformational states, are modeled as occurring at the end of each data acquisition period and are described using transition probabilities. Yet, while measurements are often performed at discrete time levels in the Natural Sciences, physical systems evolve in continuous time according to transition rates. It then follows that the modeling assumptions underlying the HMM are justified if the transition rates of a physical process from state to state are small as compared to the data acquisition rate. In other words, HMMs apply to slow kinetics. The problem is, as the transition rates are unknown in principle, it is unclear, *a priori*, whether the HMM applies to a particular system. For this reason, we must generalize HMMs for physical systems, such as single molecules, as these switch between discrete states in *continuous time*. We do so by exploiting recent mathematical tools developed in the context of inferring Markov jump processes and propose the hidden Markov jump process (HMJP). We explicitly show in what limit the HMJP reduces to the HMM. Resolving the discrete time discrepancy of the HMM has clear implications: we no longer need to assume that processes, such as molecular events, must occur on timescales slower than data acquisition and can learn transition rates even if these are on the same timescale or otherwise exceed data acquisition rates.

## 1 Introduction

Hidden Markov models (HMMs) have been important tools of time series analysis for over half a century (1–3). Under some modeling assumptions, detailed shortly, HMMs have been used to self-consistently determine dynamics of physical systems under noise and the properties of the noise obscuring the system’s dynamics itself.

Originally developed for applications in speech recognition (4, 5), the relevance of HMMs to single molecule time series analysis was quickly recognized (6–14). Since then, HMMs and variants have successfully been used in the interpretation of single ion channel patch clamp data (15–19); fluorescence resonant energy transfer (FRET) (20–28); force spectroscopy (29–31); amongst many other physical applications (4, 32, 33).

In order for HMMs to apply to single molecules and other physical systems, the assumptions underlying the HMMs must hold for such systems. There are several such assumptions worthy of consideration.

- HMMs assume that the system under study evolves in a discrete state space (whether physical of conformational). This is a reasonable approximation for biomolecules visiting different conformations (33–35) or fluorophores visiting different photo-states (33, 34, 36). Of parallel interest to this point is the notion that the number of discrete states is known (though the transition probabilities between states is unknown). The assumption of a known number of states has been lifted thanks to extensions of the HMM (34, 37–43) afforded by nonparametrics that we discuss elsewhere (33, 34, 38–40, 43).
- HMMs assume that measurements are obtained at discrete time levels. That is, successive measurements are reported at regular time levels separated by some fixed period Δ*t*. For clarity, we call Δ*t* the *data acquisition period*. This assumption is consistent with a number of experimental biophysical settings (11, 12, 38–40, 44–56).
- HMMs assume that physical systems transition between states in discrete time steps. Put differently, HMMs apply under the implicit assumption that the underlying system switches between states rarely as compared to the data acquisition period, Δ*t*. This can only really be assured if the transition rates (as required in a continuous time description) are slow. This assumption is implicit to the very definition of a HMM which requires that the system’s switching occurs *precisely* at the end of the data acquisition periods (4, 38, 39, 57–59).

This last assumption is problematic and presents the following conundrum: on the one hand, the transition rates are unknown and their analog for discrete time processes (transition probabilities) are typically to be determined using HMMs. On the other hand, we must assume that these unknowns are slow as compared to the data acquisition rate. Even if, optimistically, transition rates are slow, molecular events themselves are stochastic and, as with all physical processes, occur in continuous time. As such, any one event has a probability of occurring on timescales faster than the data acquisition period.

As an example, Fig. 1 illustrates the types of dynamical measurements that we can and cannot analyze within the HMM paradigm. The top panel shows an example of single molecule measurements characterized by slow kinetics that can be analyzed within an HMM paradigm. In contradistinction to the above is the bottom panel illustrating an example of fast kinetics, as compared to the data acquisition rate, that cannot currently be analyzed within the HMM paradigm. The reason for this is simple: the fast kinetics give rise to a large number of apparent states that go beyond the two real states. This is because the measurements reported at each time point averages the molecular signal from all states visited in each acquisition period. Yet it is clear that the information on the transition rates between the rapidly switching molecular states is encoded in the time trace however uninterpretable.

**Fig. 1.**
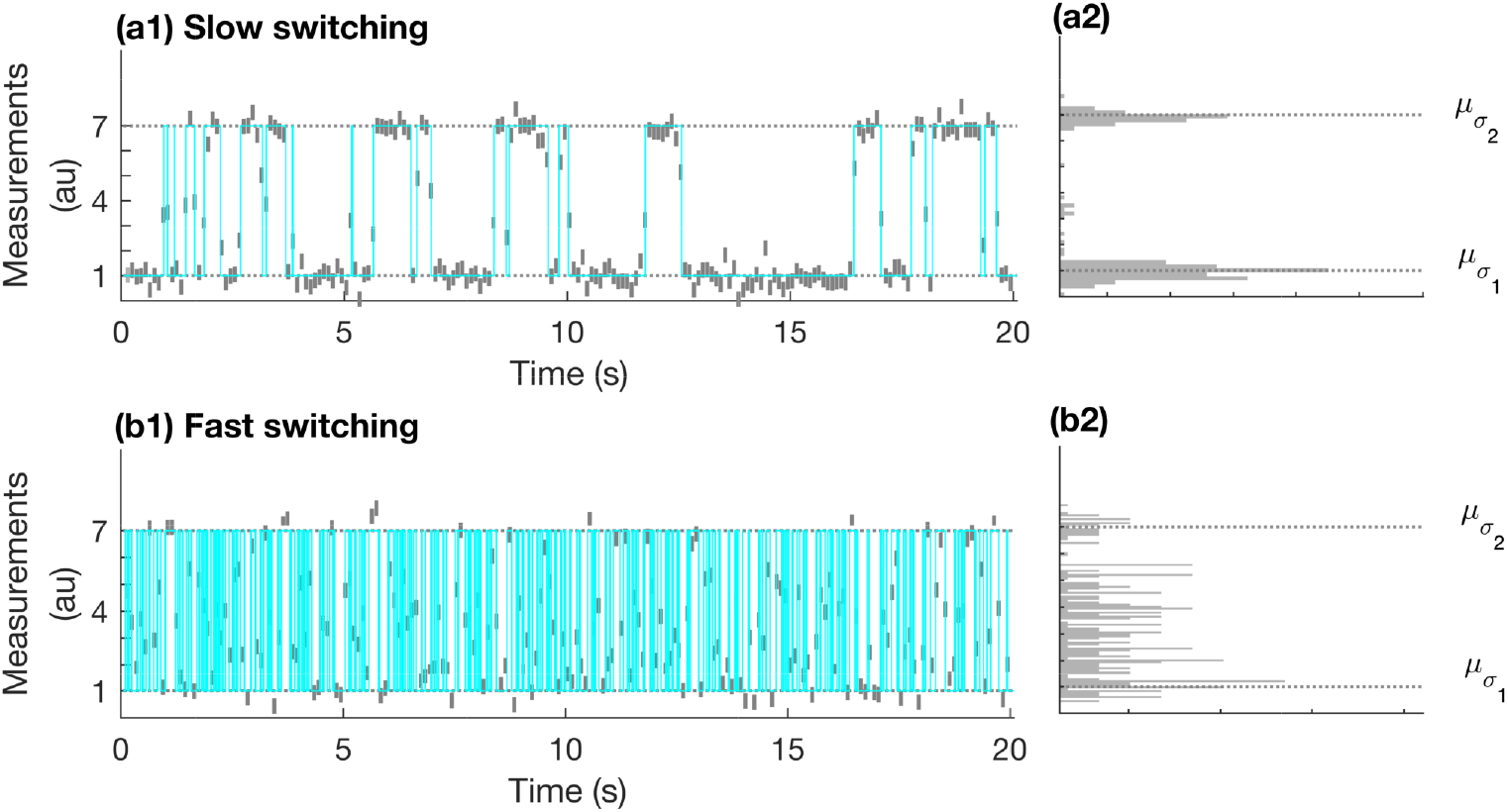
A conceptual illustration of single molecule continuous time switching kinetics between discrete states probed in discrete time. For illustrative purposes only, the trajectory of a single molecule between two states (σ_1_, σ_2_) is shown in cyan in panel (a1) and (b1). For concreteness, we can think of these molecular states as conformational states. The state levels, i.e., signal level in the absence of noise, for these states is μ_σ_1__ and μ_σ_2__, respectively, and shown by the horizontal gray dotted line in (a1) and (b1). This synthetic experiment starts at time t_0_ = 0.05 s, and ends at time t_N_ = 20 s and the data acquisition period, Δt, is 0.1 s. Next, again only for illustration, we assume that the measurements are acquired by a detector that has a fixed integration period τ = 90 ms (light gray) for each fixed data acquisition period shown in panels (a1) and (a2). As a molecule may switch between states during an integration period, the measurements represent the signal levels capturing the average of the amount of time spent at each state levels (μ_σ_1__, μ_σ_2__) visited (in addition to added noise). Panels (a1)-(a2) are associated with single molecule kinetics that are slower than the data acquisition rate. Instead, in panels (b1)-(b2) we show a single molecule trajectory when a molecule’s kinetics are faster than the data acquisition rate. In panel (b1), slow kinetics result in well separated state occupancy histograms around the average state levels. In panel (b2), we don’t have well separated histograms centered around the average state levels due to fast kinetics of the molecule.

Indeed, to address the last assumption, a recent method (45), termed H^2^MM, was proposed. H^2^MMs have been applied to single molecule FRET (smFRET) photon arrival time series analyses (46, 47). They handle fast switching kinetics within an HMM framework by embedding a finer discrete timescale into the HMM; in this case one fine enough to avoid the arrival of two photons in one discrete time bin. The H^2^MM applies to a scenario different from that provided in Fig. 1 for which the detector model produces measurement that coincide with noise on top of average molecular signal obtained throughout the detector’s exposure time.

Statistical analysis methods exploiting finer time grids, to approximate faster *continuous* time processes, had previously been considered albeit for applications outside the Natural Sciences (60, 61). Such statistical methods (60, 61) have been criticized (62) for two main reasons: 1) they sometimes, though not always, introduce computational complexity associated with a finer time grid; and, almost certainly, 2) introduce bias by discretely approximating a continuous time process. In the mathematical literature, these two challenges are what motivated the development of strategies to infer kinetic rates for genuinely continuous time processes albeit measured in discrete time (62).

It is therefore natural for us to propose an analysis method that treats physical processes as they occur in *continuous* time in order to extract rates directly from traces with fast kinetics without relying on the artificial assumption that the physical processes involved occur on timescales much slower than the data acquisition period.

To do so, we must fundamentally upgrade both key ingredients of the HMM model: 1) the system dynamics must be in continuous time; and 2) the measurement output must realistically reflect an average over the dynamics of the system over the data acquisition period. The output is then understood to encode fast dynamics, that can be retrieved (56, 63–67).

It is indeed to address processes evolving in continuous time that continuous time Markov models, so-called Markov jump processes (MJPs), were developed (44, 68–70). MJPs describe continuous time events using rates (rather than transition probabilities) and recent advances in computational statistics (62, 71–75), have made it possible to learn these rates given data. However, an important challenge remains for us to infer MJP rates under the assumption that the measurement process averages the probed signal over each measurement period. The nature of the measurement process and continuous dynamics therefore suggest a Hidden MJPs (HMJPs) framework that we put forward herewith.

In Section 2, below, we start with the formulation of our HMJP model and also, briefly, summarize the HMM. Next, in Section 3, we move on to the head-to-head comparison of HMMs and HMJPs (showing in what limit the HMM exactly reduces to the HMJP). We focus on their respective performance in learning molecular trajectories and transition probabilities. We show how HMJPs successfully outperform HMMs especially for kinetics occurring on timescales on the order of or exceeding the data acquisition period. Finally, in Section 4, we discuss the broader potential of HMJPs to Biophysics. Fine details on the implementation of these two methods can be found in Supporting Material (A).

## 2 Methods

In this section we describe a physical system that evolves in continuous time alongside a measurement model. We also discuss how to generate realistic synthetic data from such a model and subsequently analyze time traces reflecting both fast and slow dynamics. We analyze the traces using two different methods: HMMs, as they are broadly used across the literature, and our proposed HMJPs which we describe in detail. We compare the analyses in Section 3.

### 2.1 Model Description

Using the experimental data, and the model of the experiment that we will describe, our goal is to learn: 1) the switching rates between the states of the system; 2) the state of the system at any given time (that we call the trajectory of the system); 3) initial conditions of the system; and 4) parameters describing the measurement process, i.e., parameters of the emission distribution.

#### 2.1.1 Dynamics

We start by defining the trajectory 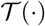 that tracks the state of the system over time. Here 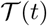 is the state of the system at time *t* and, as such, it is a *function* over the time interval [*t*_0_, *t_N_*]. We adopt functional notation and distinguish between 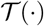 and 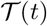 to avoid confusion with the entire trajectory and the value attained at particular time levels, critical to the ensuing presentation.

We label the states to which the system has access with *σ_k_* and use the subscript *k* = 1,…, *K* to distinguish them. For example, *σ*_1_/*σ*_2_ may represent a protein in folded/unfolded conformation or an ion channel in on/off state. With this convention (borrowed from (62)) if the system is at *σ_k_* at time *t*, then we write 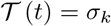. An early point of distinction with the HMM is warranted: the switching times between states of the system, generally, differ from the measurement acquisition levels *t_n_*.

As with most molecular systems (35, 38, 76, 77), the switching dynamics are faithfully modeled as memoryless. That is, the waiting time in a state of the system is exponentially distributed. Such memoryless systems are termed *Markov jump process* and below we present their mathematical formulation.

At the experiment’s onset, we assume the state of the system 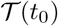 is chosen stochastically among *σ_k_*. We use *ρ_σ_k__* to denote the probability of the system starting at *σ_k_* and collect all initial probabilities in 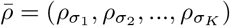, which is a probability vector (78–81).

Memoryless switching kinetics are described by *switching rates* between all possible state pairs. These switching rates are labeled with *λ_σ_k_→σ_k′__*. By definition, all self-switching rates are zero *λ_σ_k_→σ_k__* = 0, which, in general, allows for at most *K*(*K* – 1) non-zero rates (35). Although, the switching rates *λ_σ_k_→σ_k′__* fully describe the system’s kinetics, as we will see shortly, it is mathematically more convenient to work with an alternative parametrization.

In this alternative parametrization, we keep track of the *escape rates*

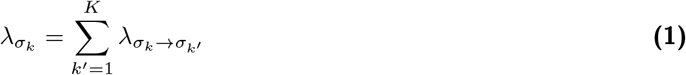

which, for simplicity, we gather in 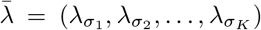. Further, instead of keeping track of each rate *λ_σ_k_→σ_k′__*, we keep track of the rates normalized by the escape rates, namely

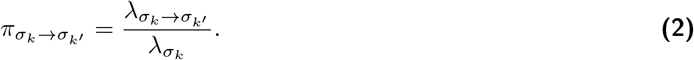

Gathering all normalized rates out of the same state in 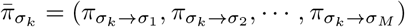, we see that each 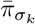 forms a probability vector (80).

In summary, instead of *K*(*K* – 1) switching rates *λ_σ_k_→σ_k′__*, we describe the system’s kinetics with *K* escape rates *λ_σ_k__* and *K* switching probability vectors 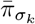. The latter have, by definition, *π_σ_k_→σ_k__* = 0, and so, the total number of scalar parameters is the same in both parametrizations. Below, for simplicity, we gather all transition probability vectors into a matrix

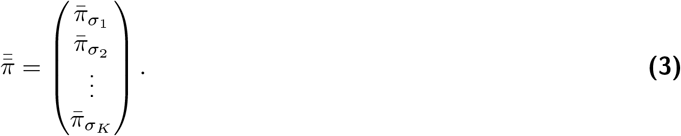

#### 2.1.2 Measurements

The overall input to our method consists of the measurements **x** = (*x*_1_, *x*_2_,…, *x_N_*) acquired in an experiment. Here, *x_n_* indicates the *n*^th^ measurement and, for clarity, we assume measurements are time ordered, so *n* = 1 labels the earliest acquired measurement and *n* = *N* the latest. These measurements may be camera ADUs generated from photon detections, FRET efficiencies, derived inter-molecular extensions, or any other quantity determinable in an experiment.

Each *x_n_* is reported at a time *t_n_* = *t*_*n*–1_ + Δ*t*, which is Δ*t* later than the time *t*_*n*–1_ at which the previous measurement *x*_*n*–1_ was reported. For completeness, together with the time levels *t*_1_, *t*_2_,…, *t_N_* at which a measurement is reported, we also consider an additional time level *t*_0_, that marks the onset of the experiment, which is not associated with any measurement, Fig. 1.

The most common assumption made almost universally by HMMs is that the instantaneous state of the system at *t_n_* determines *x_n_*. Yet, for realistic detectors, the reported value *x_n_* is influenced by the entire trajectory of our system during the *n*^th^ integration period which we represent by the time window [*t_n_* – *τ,t_n_*]. Here, *τ* is the duration of each integration time (such as an exposure period for optical experiments).

We account for detector features in the generation of the measurements via characteristic state levels which we label with *μ_σ_k__* and, for simplicity, gather these in 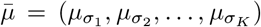. In our formulation, each *σ_k_* is associated with its own characteristic level *μ_σ_k__*. If the system remains at a single state *σ_k_* throughout an entire integration period [*t_n_* – *τ, t_n_*], then the detector is triggered by *μ_σ_k__* and so, provided measurement noise is negligible, the reported measurement *x_n_* is similar to *μ_σ_k__τ*. However, if the system switches multiple states *during* an integration period, the detector is influenced by the levels of every state attained and the time spent in each state.

More specifically, the *n*^th^ signal level triggering the detector during the *n*^th^ integration period, [*t_n_* – *τ,t_n_*], is obtained from the time average of 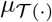 over this integration period. Mathematically, this time average equals 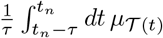 and, provided measurement noise is negligible, the reported measurement *x_n_* is similar to the value of this average.

In the presence of measurement noise, such as shot-noise (82–86), quantification noise (87–89), or other degrading effects common to detectors currently available, each measurement *x_n_* depends *stochastically* upon the signal that triggers the detector (33, 90, 91). Of course, the precise relationship depends on the detector employed in the experiment and differs between the various types of cameras, single photon detectors or other devices used. To continue with our formulation, we assume that measurement noise is additive, which results in

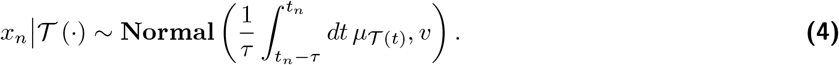

The latter expression is a statistical shorthand for the following: the measurement *x_n_* is a random variable that is sampled from a normal distribution whose mean is 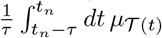 and whose variance is *v*. For the Normal distribution, the variance is related to the detector’s full-with-at-half-maximum (FWHM) by 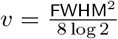.

Our choice of a Normal distribution itself is incidental (and can be changed depending on detector type). However, this type of measurement model is general enough to capture the effect of the history of the system during the detector’s integration time. For example, in an accompanying article (92), we adapt Eq. (4) to FRET measurements in separate donor and acceptor channels with shot-noise and background as follows

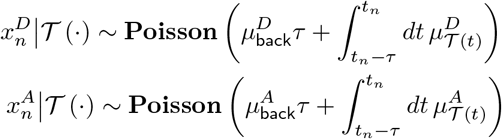

where 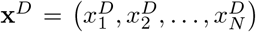 and 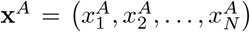 denote the measurements acquired in the donor’s and the acceptor’s channels, respectively. Here, 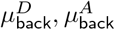 denote the background and 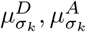 the characteristic state levels in the two channels.

#### 2.1.3 Simulation

Given 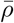 and 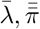 a trajectory 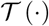 that mimics real systems may be simulated using the Gillespie algorithm (76) which we describe briefly here only in an effort to introduce necessary notation.

To begin, an initial state *s*_0_ is chosen among *σ*_1_, *σ*_2_, ···, *σ_K_* with probability *ρ_σ_k__*. Then, the period *d*_1_ that the system spends in *s*_0_ is chosen from the Exponential distribution with mean 1/λ_*s*_0__. Subsequently, the next state *s*_1_ is chosen among *σ*_1_, *σ*_2_, ···, *σ_K_* with probability *π*_*s*_0_→*σ_k_*_. As *π*_*s*_0_→*s*_0__ = 0, any chosen *s*_1_ is different from *s*_0_, therefore the transition *s*_0_ → *s*_1_ is a jump in the system’s time course that occurs at time *t*_0_ + *d*_1_. Next, a new period *d*_2_ is sampled from an Exponential distribution with mean 1|λ_*s*_1__ and a new state *s*_2_ is chosen among *σ_k_* with probability *π*_*s*_1_→*σ_k_*_, and so on. These steps are repeated until the end of the experiment which, in our setup, is the same as the time *t_N_* of the last measurement.

More formally, we summarize the sampling of a Gillespie trajectory as follows

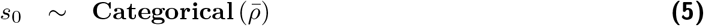

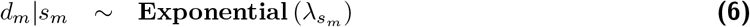

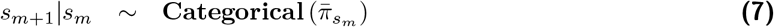

for *m* = 0, 1, 2,…, *M* – 1 where *M* – 1 is the lowest value such that

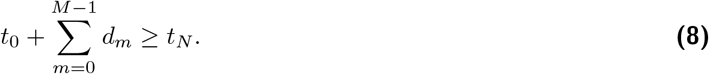

The Categorical distribution we use here is the generalization of the Bernoulli distribution for which more than two outcomes are possible (57).

The successive states of the system *s*_0_, *s*_1_, ···, *s*_*M*–1_ and the associated durations *d*_0_, *d*_1_, *d*_2_, ···, *d*_*M*–1_, which we term *holding states* and *holding times*, respectively, encode 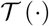 throughout the experiment’s time course [*t*_0_, *t_N_*]. Namely

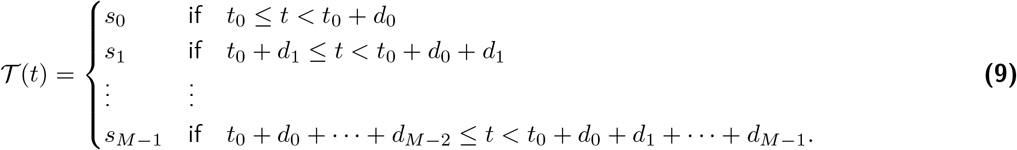

For convenience, we summarize the representation of 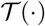 in a triplet 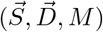, where

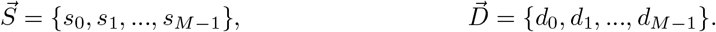

and *M* is the size of 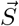 and 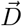.

Once a trajectory is obtained through the Gillespie algorithm just described, the signal levels 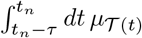 for each integration period can be computed. For instance, as the trajectory is piecewise constant, the integrals reduce to sums that can be easily calculated. Therefore, given an appropriate detector model, such as Eq. (4), and a trajectory’s triplet 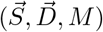, we can obtain simulated measurements by adding noise according to the detector’s distribution.

### 2.2 Model Inference

Using the data **x**, and the model of the experiment that we have just described, our goal is now to learn initial probabilities *ρ_σ_k__*, switching rates *λ_σ_k_→σ_k′__*, and state levels *μ_σ_k__* for all states as well as the trajectory of the system 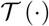 throughout the experiment’s time course [*t*_0_, *t_N_*]. Below we attempt to learn these using time series analysis with a HMM and then introduce a novel time series analysis relying on HMJPs.

#### 2.2.1 Model Inference via HMMs

An HMM requires that each measurement *x_n_* depends exclusively on the *instantaneous* state of the system, namely 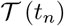. In view of Eq. (4), this is achieved by the trajectory of the system 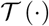 remaining constant during the integration period [*t_n_* – *τ, t_n_*]. To a sufficiently good approximation, this is satisfied provided

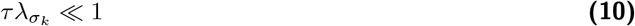

for all *σ_k_*. Thereby, the system rarely exhibits switching during periods that last shorter than *τ*. This approximation allows for 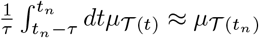 to be used. Accordingly, in a HMM, Eq. (4) is replaced with

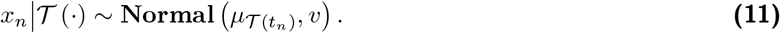

Again, as with Eq. (4), the exact choice of distribution (whether Normal or otherwise) is incidental. HMMs can treat any emission distribution *provided x_n_* only depends on 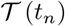 as opposed to the full history of the trajectory over the integration time.

With the measurements described by Eq. (11), we can use a HMM to learn the probabilities of the transitions 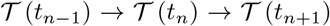. For clarity, from now on, we will use 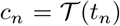 and denote these transitions with *c*_*n*–1_ → *c_n_* → *c*_*n*+1_. That is, *c_n_* is the state of the system *precisely* at the time *t_n_*.

For an HMM, transition probabilities are denoted with *P*_*c*_*n*–1_→*c_n_*_. Since the system can attain *K* different states *σ_k_*, in general, a HMM possesses *K* × *K* transition probabilities *P_σ_k_→σ_k′__*. Now, we gather the transition probabilities out of the same *σ_k_* in a vector 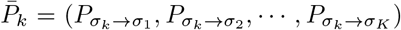 and, for clarity, gather all of the vectors in a matrix

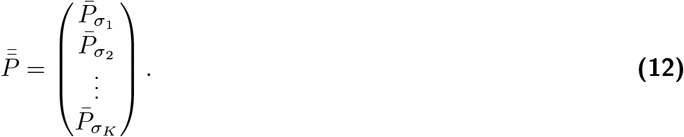

The matrix 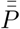 is related to the system’s switching rates *λ_σ_k_→σ_k′__* and escape rates *λ_σ_k__*. Specifically, if we gather them in

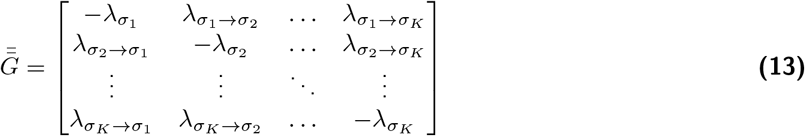

which is termed *generator matrix* (80, 93), then 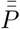 is obtained by

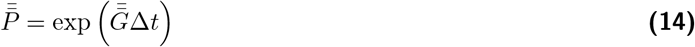

where exp(·) denotes the matrix exponential. We point out that 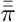 and 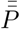 are both probability matrices, however, they assume quite *different* properties. For instance, *π_σ_k_→σ_k__* = 0 while *P_σ_k_→σ_k__* > 0.

Although knowing 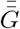 is sufficient to specify 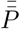, the inverse is *not* true: knowing 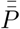 does not necessarily lead us to a unique 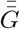 and so the switching rates *cannot* simply be inferred from 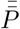. This is a consequence of the multivalued nature of the logarithm. As such, one transition probability matrix may corresponds to multiple rate matrices (93). Instead, provided *λ_σ_k__*Δ*t* ≪ 1 for all *σ_k_*, we may approximate Eq. (14) by

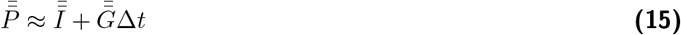

where 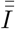 is the identity matrix of size *K × K*. Under this approximation, we can estimate transition rates by 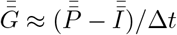.

Below, we highlight the steps necessary to estimate the quantities of interest in a HMM. Specifically, a HMM relies on the statistical model

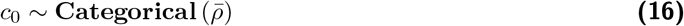

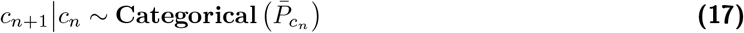

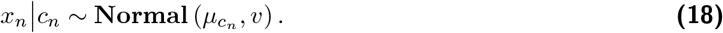

To model the full distribution over the quantities of interest, e.g., initial probabilities, 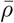, transition probabilities 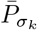, state levels 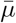 and the trajectory of the system 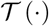 which is encoded by 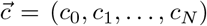, we follow the *Bayesian paradigm* (78, 94). Within this paradigm, we place prior distributions over the parameters and we discuss the appropriate choices next.

On the transition probabilities 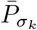, we place a Dirichlet prior with concentration parameter *A*

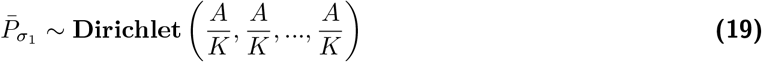

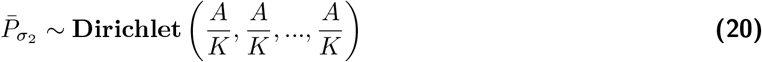

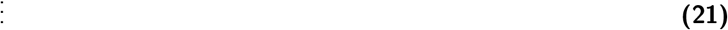

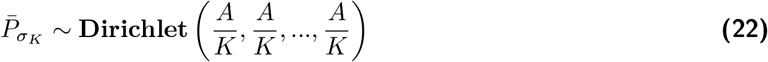

which is conjugate to the Categorical distribution (38, 39, 41, 81). We consider a similar prior distribution, with concentration parameter *α*, also for the initial transition probability 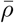, namely

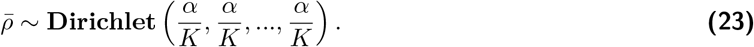

Subsequently, we place priors on the state levels 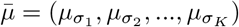. The prior that we choose is the conjugate Normal prior

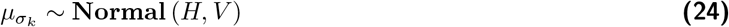

with hyperparameters *H, V*.

Once the choices for the priors are made, we then form the posterior distribution (34, 38–43)

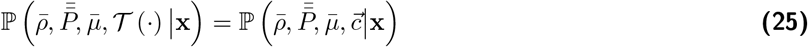

containing all unknown variables that we wish to learn. As the posterior distribution in Eq. (25) does not attain an analytical form, we develop a specialized computational scheme exploiting Markov Chain Monte Carlo (MCMC) to generate pseudorandom samples from this posterior. We explain the details of this scheme in Section 2.2.3.

#### 2.2.2 Model Inference via HMJPs

HMJP apply directly on the formulation of Section 2.1.2 and, unlike with HMM (see Eq. (10)), no approximations are required on the system switching kinetics. Therefore, in order to proceed with inference, we need only provide appropriate prior distributions on the parameters, namely 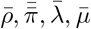.

We start with the prior distribution for the escape rates 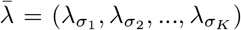. We put priors on each of the *λ_σ_k__* for all *k* = 1, 2,…, *K*. The prior we select is

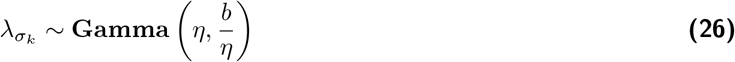

for all *k* =1, 2,…, *K* with hyperparameters *η, b*. We note that this prior is conjugate to the exponential distribution given in Eq. (6). Next, we place a prior on 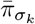 for all *k* = 1, 2,…, *K*. For this, we place independent conjugate Dirichlet priors with concentration parameter *A* such that *π_σ_k_→σ_k__* = 0 holds for all *k* = 1, 2,…, *K*

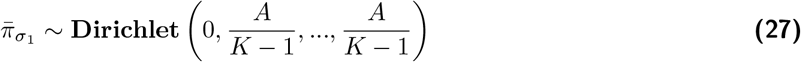

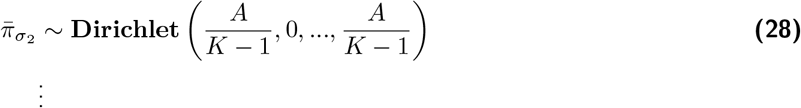

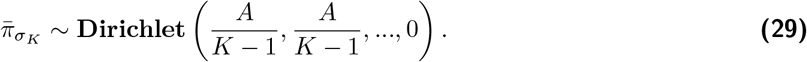

Finally, on 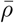 and 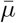 we place the same prior distributions as in Eq. (23) and Eq. (24), respectively.

Once the choices for the priors are made, we then form the posterior distribution

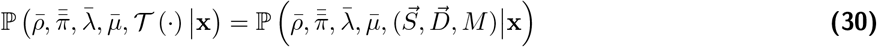

containing all unknown variables that we wish to learn. As, once more, the posterior distribution does not attain an analytical form, we develop a specialized computational scheme exploiting MCMC. We explain the details of this scheme in Section 2.2.3.

#### 2.2.3 Computational Inference

We carry out the analyses, shown in Section 3, evaluating the associated posteriors with an MCMC scheme relying on Gibbs sampling (34, 38–43, 62). The overall sampling strategy, for either the HMM or the HMJP, is as follows:

1. Update the trajectory 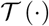, that is 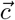 for HMM or 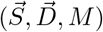 for HMJP;
2. Update the kinetics, that is 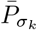 for HMM or 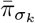 and *λ_σ_k__* for HMJP;
3. Update the initial probabilities 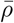;
4. Update state levels 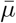.

We repeat these updates to obtain a large number of samples. The end result is a sampling of the posterior 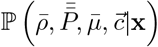 for the HMM and 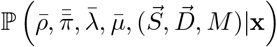 for the HMJP. Both samplings can be used to estimate switching rates *λ_σ_k_→σ_k′__*; for example, HMJP by Eq. (2) and HMM by Eq. (15).

In the Supporting Material (A.2) we provide a thorough description of all steps. We also freely provide a working code through the authors’ website.

## 3 Results

To demonstrate how HMJPs work and highlight their advantages over HMMs, in this section, we use synthetic data that mimic a single molecule experiment. Synthetic data are ideal for this purpose as they allow us to benchmark the results against the exact, readily available, “ground truth”. We obtain such data from the Gillespie algorithm described in Section 2.1.3 and we explain our simulation choices below.

We focus on two datasets: one where the system exhibits slow kinetics and another where the system exhibits fast kinetics as compared with data acquisition, see Fig. 1. We provide the values for the hyperparameters in all analyses, as well as any other choices made, in Supporting Material (A.3). To be clear, we only assume to have access to the data, i.e., the gray dashes of panels (a1) and (b1) of Fig. 1. The cyan (ground truth) trajectories are assumed unknown and to be determined.

In our results, we first benchmark the HMJP on the easy (i.e., slow kinetics) case shown in Fig. 1 panel (a1); see Figs. 2 and 3. This is the regime where the HMM also works well and the expected (good) results for the HMM are relegated to the appendix; see Supporting Material (A.1). Next, we turn to the more complex case of fast kinetics. A sample time trace is shown in Fig. 1 panel (b1). The results for both the HMJP and HMM are shown in Figs. 4 and 5.

**Fig. 2.**
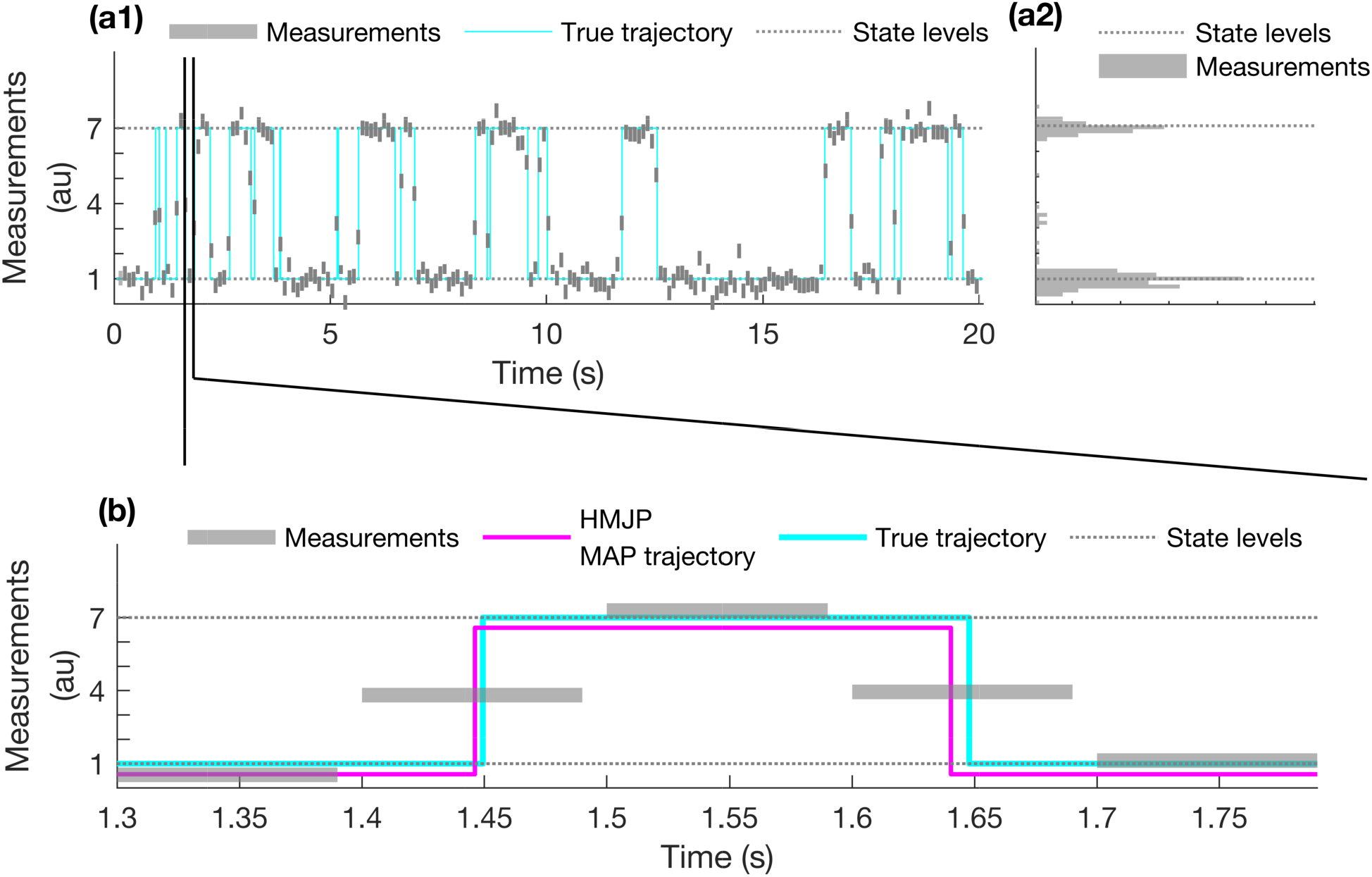
HMJP trajectory estimates for slow state switching. Here we provide trajectory estimates obtained with the HMJP when the switching rate is slower than the data acquisition rate, 1/Δt = 10 (1/s). In this figure’s panel (a1), the measurements are shown as gray rectangles (the width of the rectangle coincides with the integration period as shown in Fig. 1) generated based on the description provided in Section 2.1. We superposed the true trajectory (cyan) with the measurements in panel (a). Next, in panel (a2), we provide the histogram of all measurements to visualize the system kinetics. For illustrative purposes, we only show the MAP estimates of the HMJP on a zoomed in region of panel (a1). Next, we provide that region of the panel (a1) in panel (b). In panel (b), we show the the MAP trajectory estimates of HMJP (magenta) that is superposed with the measurements and the true trajectory (cyan). For visual purposes only, we offset the HMJP MAP trajectory estimate by slightly shifting it downward. We observe that the HMJP MAP trajectory is able to capture switching occurring roughly in the middle of the integration time. This is not something that the HMM can capture. Here, simulated measurements are generated with λ_σ_1_→σ_2__, λ_σ_2_→σ_1__ Eq. (31) where the data acquisition happens at every Δt = 0.1 s with τ_f_ = 0.8 s and τ = 0.09 s starting at t_0_ = 0.05 s until t_N_ = 20 s.

**Fig. 3.**
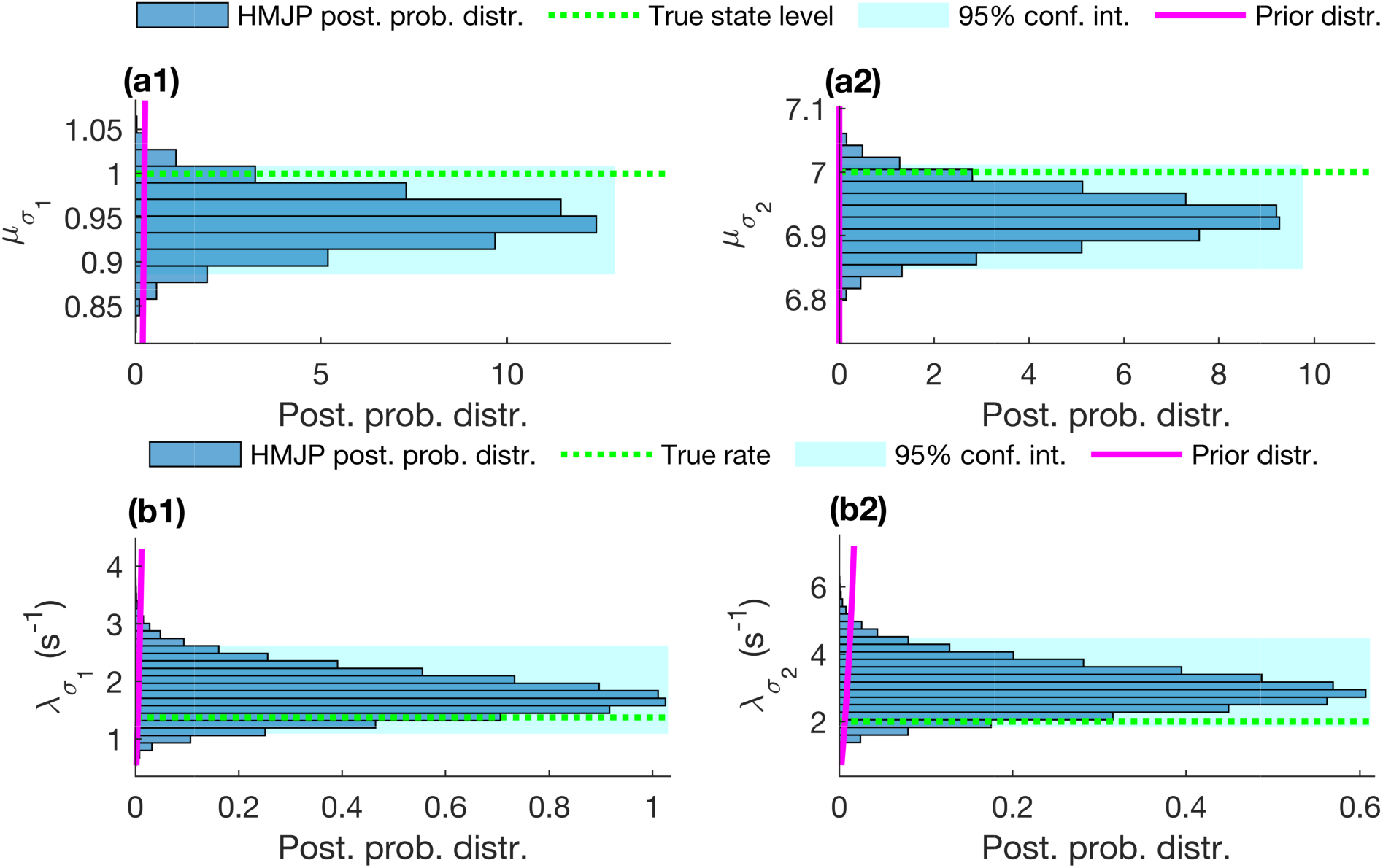
HMJP state level and rate estimates for slow state switching. Here we provide posterior state level and rate estimates obtained with HMJP whose time trace we discussed in Fig. 2. We expect HMJPs to perform well in estimating the true state levels and rates when these rates are slower than the data acquisition rate. In all figure panels, we superposed the posterior distributions over state levels and rates for HMJP (blue) along with their 95% confidence intervals, the true state levels (dashed green lines) and the corresponding prior distributions (magenta lines). We start with the information in panels (a1) and (a2). We observe in these panels that the HMJP posterior distributions over state levels contain the true state levels within their 95% confidence intervals. Next we move to the panels (b1) and (b2) which show the posterior distributions over the rates labeled λ_σ_k_→σ_k′__ for all k, k′ = 1, 2. Again, the HMJP does quite well in estimating these rates as measured by the fact that the ground truth lies within the 95% confidence intervals of the posteriors. In this figure, the analyzed simulated measurements are generated with the same parameters as those provided in Fig. 2.

**Fig. 4.**
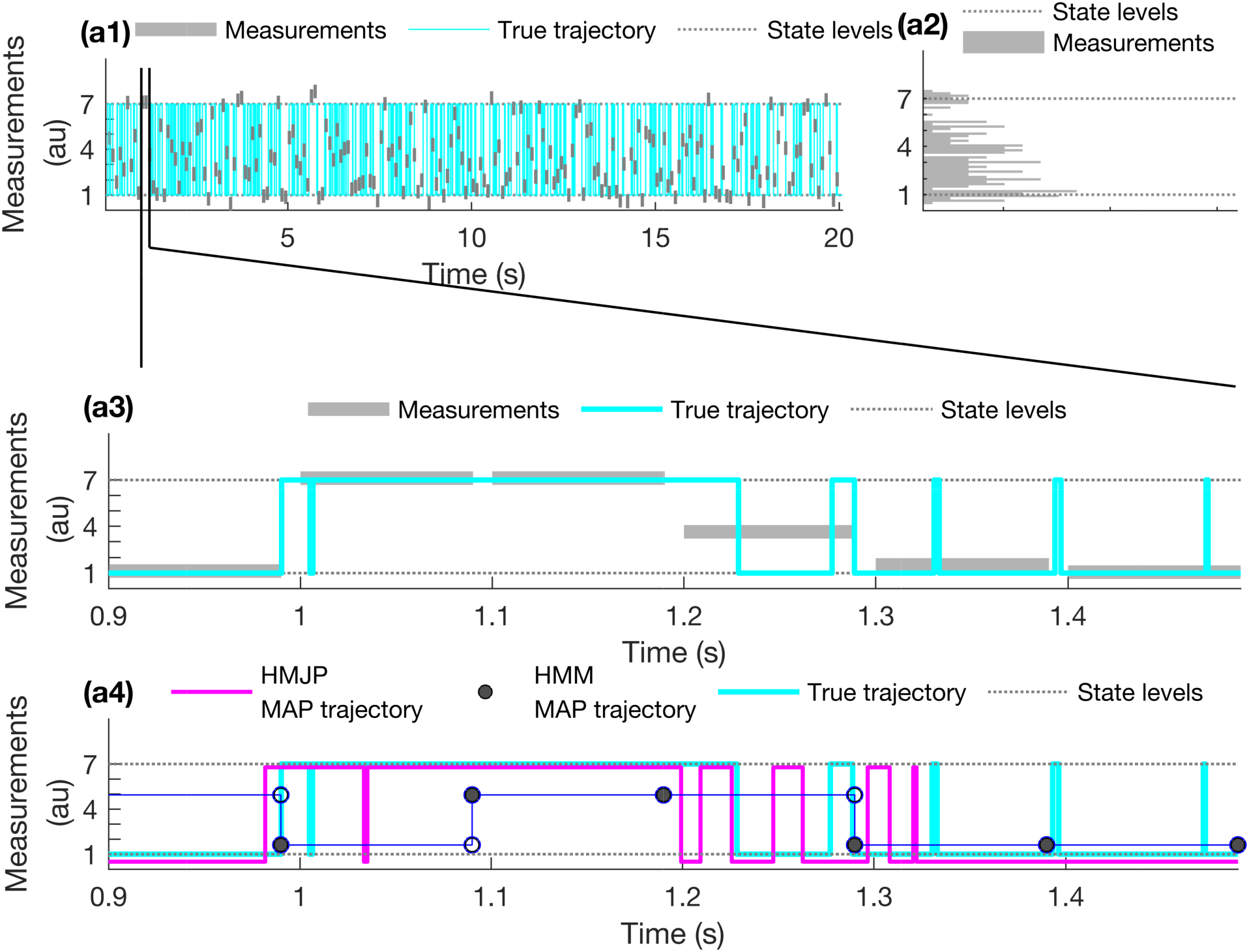
HMJP with HMM trajectory estimates for fast state switching. Here we provide trajectory estimates obtained with HMJP and HMM when the switching rate is faster than the data acquisition rate, 1/Δt = 10 (1/s). We expect HMMs to perform poorly in estimating the true trajectory when switching is fast. In this figure’s panel (a1), the measurements are shown with gray rectangles (the width of the rectangle coincides with the integration period as shown in Fig. 1) that are generated based on the description provided in Section 2.1. We follow the same color scheme and layout as in Fig. 2 except for panel (a4) where we provide the MAP trajectory estimate provided by the HMJP as well as the HMM. In panel (a4), the magenta dashed line shows the HMJP MAP trajectory estimate and the blue line shows the HMM MAP trajectory estimate. For visual purposes, we offset the HMJP MAP trajectory estimate and HMM MAP trajectory estimate by shifting these downward. Here, simulated measurements are generated with λ_σ_1_→σ_2__, λ_σ_2_→σ_1__ (see Eq. (31)) where the data acquisition happens at every Δt = 0.1 s with τ_f_ = 1/15 s and τ = 0.09 s starting at t_0_ = 0.05 s until t_N_ = 20 s.

**Fig. 5.**
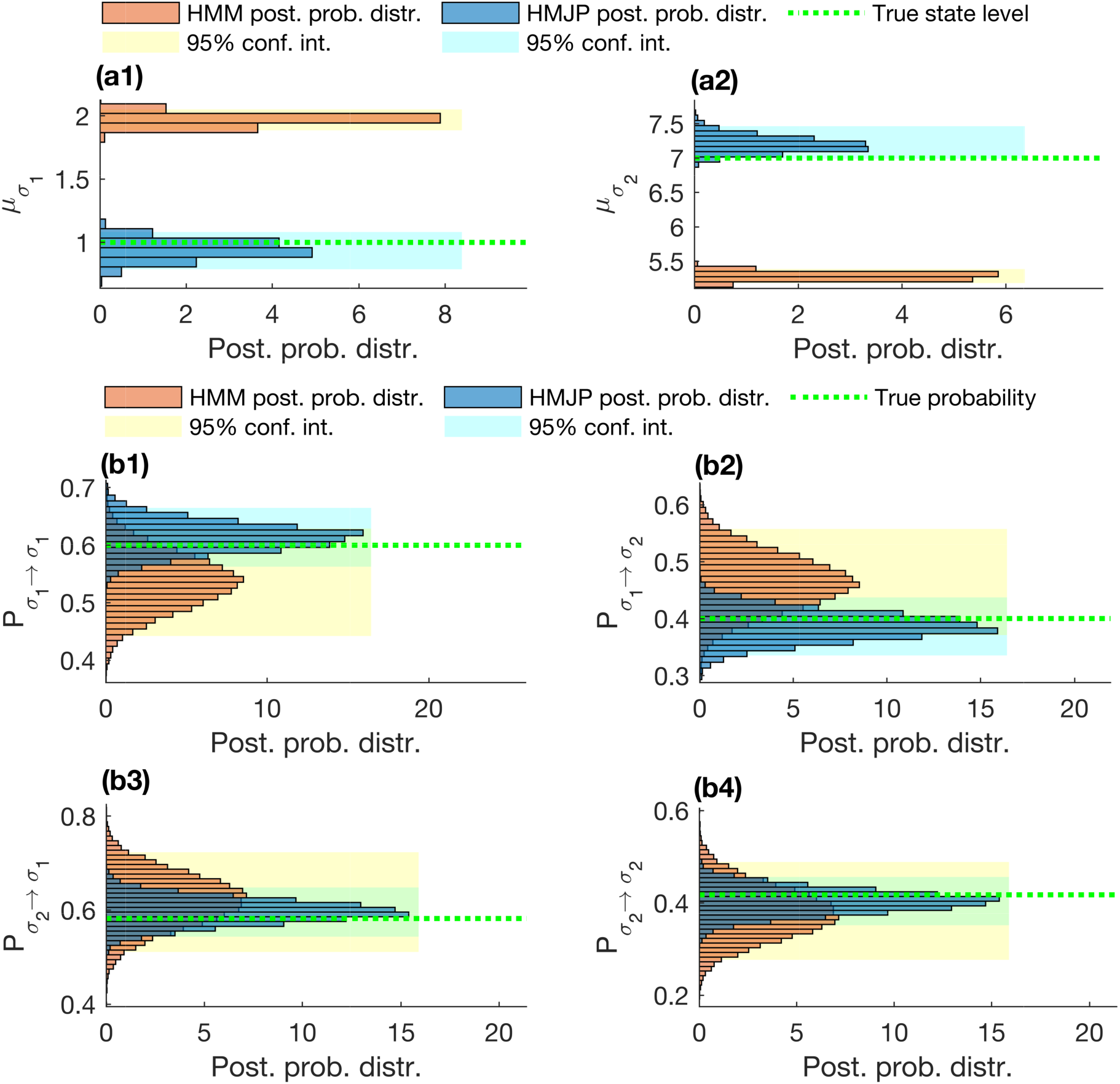
HMJP with HMM state level and transition probability estimates for fast state switching. Here we provide posterior state level and transition probability estimates obtained with HMJP and HMM when the switching rate is faster than the data acquisition rate, 1/Δt = 10 (1/s). We expect HMMs to perform poorly in estimating the true state levels and transition probabilities when the system switching is fast. In all of this figure’s panels, we superposed the posterior distributions over state levels for both HMJP (blue) and HMM (orange) along with their 95% confidence intervals, the true state levels and true transition probabilities (dashed green lines). Next we move to transition probability estimates provided in panels (b1)-(b4). In these panels, we wish to test the performance of HMJPs and HMMs in estimating the transition probabilities. In this figure, each panel is corresponding to the posterior distribution of a transition probability labeled as P_σ_k_→σ_k′__ for all k, k′ = 1, 2. Here, simulated measurements are generated with the same parameters as those provided in Fig. 4.

### 3.1 Data Simulation

To simulate the synthetic data, we assumed *K* = 2 distinct states, such as on/off or folded/unfolded states for illustrative purposes only. We assumed well separated state levels which we set at *μ*_*σ*_1__ = 1 au and *μ*_*σ*_2__ =7 au where au denote arbitrary units. The prescribed detector FWHM was set at 0.25 au.

Additionally, for sake of concreteness only, we assumed an acquisition period of Δ*t* = 0.1 s and consider long integration periods by setting *τ* equal to 90% of Δ*t*. In terms familiar to microscopists, our setting corresponds to a frame rate of 10 Hz with exposure time of 90 ms and a dead time of 10 ms (40). The onset and concluding time of the experiment are the same for all simulated measurements and set at *t*_0_ = 0.05 s and *t_N_* = 20 s, respectively.

To specify kinetics, we use the following structure for the switching rates *λ*_*σ*_1_→*σ*_2__, *λ*_*σ*_2_→*σ*_1__, with a parameter *τ_f_* which sets the timescale of the system kinetics,

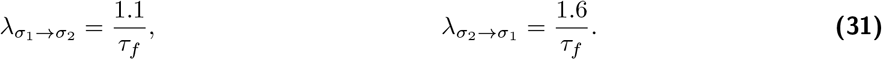

We simulate a case with *τ_f_* = 0.8 s, which involves system kinetics that are slower than the data acquisition rate; and a case with *τ_f_* = 0.067 s, which involves system kinetics that are faster than the data acquisition rate.

### 3.2 Analysis with HMJPs

As a benchmark, we provide the results for the HMJP for those measurements shown on Fig. 1 panel (a1) associated with slow switching rates. These results include estimates of the trajectory 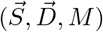 (see Fig. 2), state levels 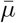 (see Fig. 3) and the switching rates 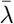 (see Fig. 3). To obtain these estimates, we generate samples from the posterior distribution 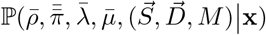 with the HMJP sampler of Section 2.2.3.

In Fig. 2 panel (a1), the ground truth trajectory is shown in cyan while the measurements are shown in gray. We showed the zoomed trajectory and observations in panel (b). We also provide the empirical histogram of the observation in panel (a2) highlighting the slow switching rates of the system. After determining the posteriors over the trajectories with HMJPs, for illustrative purposes, we only show the maximum a posteriori (MAP) trajectory in Fig. 2 panel (b). We observe that the HMJP MAP trajectory (magenta) captures most of the fast switches, shown in Fig. 2 panel (b), in the system trajectory. In Fig. 3, there are four panels. In these four panels, we provide the superposed posterior distributions over the two state levels and two rates estimated by the HMJP along with its associated 95% confidence interval and ground truths (dashed green lines).

In summary: HMJP performs well on this benchmark data. The same is true of the simpler HMM (as would be expected) whose results are shown in Figs. A.1 and A.2. An important bring home message for the HMJP however is the fact that even if state transitions occur midway through the integration time, the HMJP can discern when these occurred. The same is not true of the HMM that, as mentioned earlier, assumes by construction that state transitions must occur at the end of the data acquisition period.

### 3.3 Comparison of HMJPs with HMMs

We now present a comparison of HMJPs and HMMs on the analysis of the simulated measurements shown in Fig. 1 panel (b1). We expect the HMJP to outperform the HMM as we are now operating in a regime, with switching rates 2.5 times faster than earlier, where the HMM requirement, spelled out in Eq. (10), breaks down.

We used these measurements to estimate the posterior distribution over the trajectory, 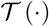, initial and switching probabilities, 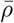 and 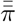 or 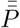, state levels, 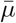 and escape rates, 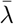. To accomplish this we generate samples from the posterior distributions 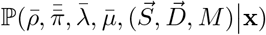 and 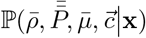 using the HMJP and HMM samplers of Section 2.2.3, respectively.

Following the pattern from the previous section on slow kinetics, we first show the trajectories inferred by HMJPs and HMMs in Fig. 4, then we show estimates of the state levels and transition probabilities in Fig. 5. In particular, the escape rates estimated from HMJPs are used in Eq. (15) to yield transition probabilities that we subsequently compared with the transition probabilities inferred by HMMs.

Predictably, the HMM performs poorly. For example, we see in Fig. 4 panel (a4) that the HMJP MAP trajectory captures many of the fast switches occurring during integration times. The HMM MAP trajectory is severely constrained to allowing switches at the end of the time period and, as such, cannot accommodate fast kinetics. While the trajectory inferred by HMJPs is not perfect, this ability to tease out many correct state switches in its MAP trajectory is sufficient for HMJPs to obtain estimates of the transition probabilities and state levels that lie within the 95% confidence interval; see panels (a1)-(a2) and (b1)-(b4) of Fig. 5. The same does not hold for HMMs where their inability to detect state switches now percolates down to the quality of their estimates for the state levels and transition probabilities. To wit, from panels (a1)-(a2) of Fig. 5 we see that the HMM grossly overestimates (by about 90%) *μ*_*σ*_1__ and underestimates (about 30%) *μ*_*σ*_2__. What is more, as can be seen in panels (b1)-(b4) of Fig. 5, the HMM provides very wide posterior distributions over transition probabilities. This is by contrast to the much sharper posterior of the HMJP whose mode closely coincides with the ground truth; see panels (b1)-(b4) of Fig. 5.

An observation is warranted here. As the HMM cannot accommodate fast kinetics, it must ascribe the apparent spread around the *P_σ_k_→σ_k′__* histogram (see panels (b1)-(b4) of Fig. 5) to an increased variance in the posterior distribution of transition probabilities. So, while the breadth of the posteriors of the HMJP are primarily ascribed to the fact that finite data informs the posterior, the origin of the breadth of the histogram of the HMM is an artifact of its inability to accommodate fast kinetics.

In the appendix, we delve into finer details of the effect of finiteness of data on the HMM posterior distributions over transition probabilities for fast switching rates in Supporting Material (A), see Fig. A.6. In particular, we analyzed a sequence of 3 data sets using the HMM framework with the same fast switching rates as in Fig. 4 panel (a1) but with differing data set lengths. While more data eventually narrows the HMM’s posterior over transition probabilities, the amount of data has almost no effect on the poor posterior distributions over state levels Fig. A.7. Further analysis on the performance of both HMJP and HMM is left for the Supporting Material (A) in particular cases where the rate from one state to another is fast and the other is slow. We also provide the HMJP rate estimates in Fig. A.5 panels (a1)-(a2) for the data set given in Fig. 1 panel (b1) as well as a comparison of HMJP posterior transition probability estimates associated with the data provided in Fig. 1 panel (a1) with and without learning the trajectory 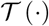 simultaneously in Fig. A.8. Finally, we compare the posterior trajectory estimates of HMJP and HMM based on a metric that is the enclosed area under the learned trajectories in Fig. A.9.

## 4 Discussion

HMMs have been a hallmark of time series analysis in single molecule Biophysics (11, 12, 38–40, 44, 45, 48–56) but they have a critical limitation. HMMs apply only provided the temporal resolution of the experimental apparatus is faster than the system kinetics under study (31, 95–97). Otherwise, HMMs mistakenly ascribe the signal generated by fast dynamics to misassignments of signal levels, such as Förster resonance energy transfer distances in single molecule FRET (7, 12, 38, 39, 48), amongst other artefacts; see Figs. 5, A.2 and A.4. Fundamentally this limitation arises because HMMs treat the dynamics of the probed systems discretely while, in fact, they evolve continuously.

The HMJP we describe here is the continuous time analog of the HMM. It can be easily adapted to treat various single molecule experiment setups. To wit, in a companion article, we will use HMJPs for the analysis of single molecule FRET (20–27, 44).

The HMJP tackles a different problem as that of the H^2^MM: 1) we work in the Bayesian paradigm and obtain full posteriors over unknowns while the H^2^MM uses maximum likelihood and provides point statistics over unknowns; 2) the HMJP tackles data from a different experiment as that of the H^2^MM. In particular, the H^2^MM assumes the data are available as single photon trajectories while we focus on the fundamental challenge of unraveling processes on timescales faster than those of detectors with finite exposure time.

The HMJP does have limitations. In the limit that state switching rate grows, the amount of data needed to ascertain a meaningful posterior over the transition kinetics also grows. In the trivial limit that the state switching is extremely fast, no method, whether HMJP or otherwise, would be able to tease out information on the transition kinetics from what appears as a uniform horizontal time trace with noise with no discernible transitions. Although, the quality of the data is not a limitation of HMJP, it is clear that the duration of the detector dead time affects the performance of all methods of inference. Specifically, the longer the dead time, the worse the HMJP will perform. In the limit the dead time is as long as the exposure itself, the HMJP reduces to the HMM.

Of great interest is the possibility to learn the number of states within an HMJP framework. That is, to re-pitch the HMJP within a Bayesian nonparametric paradigm following in the footsteps of the HMM and its nonparametric realization, the infinite HMM (37–40, 98–100). Methods have been developed to report on point statistics as they pertain to infinite MJPs (70, 101). A natural extension for us would be to fully sample from a posterior with realistic measurement models relevant to single molecule Biophysics within the nonparametric paradigm.

## ACKNOWLEDGEMENTS

SP acknowledges support from NSF CAREER grant MCB-1719537 and NIH NIGMS (R01GM134426). ASU cluster AGAVE and Saguaro are the main computational resources utilized in this study. ZK thanks Sina Jazani for his helpful suggestions on the manuscript.

## A Supporting Material

In this supplement we provide: 1) additional time series analysis to test the robustness of HMJP and HMM referenced in the main text, see Supporting Material (A.1); 2) a complete summary of the theory underlying the HMJP and HMM frameworks, see Supporting Material (A.2); 3) a summary of notational conventions and parameter choices used, see Supporting Material (A.3).

### A.1 Additional Analyses

Additional results are provided in the following order: we investigated a) the performance of HMJP and HMM in estimating the trajectory, state levels and transition probabilities on the analysis of the simulated measurements generated with slow switching rates and mixed switching rates (i.e., *λ*_*σ*_2_→*σ*_1__ fast and *λ*_*σ*_1_→*σ*_2__ slow); b) the posterior rate distributions of HMJP for fast and mixed switching cases; c) how the finiteness of the data affect HMM posterior transition probabilities; d) the source of bias in negatively skewed HMJP posterior transition probability estimates; e) the assessment of the mean of the are of posterior trajectory estimates of HMJP and HMM.

#### A.1.1 Comparison of HMJP with HMM for Slow Switching Rates

In the text in Figs. 2 and 3, we omitted the results of the slow kinetics for the HMM analysis (as this is a regime where we expect the HMM to do well). We use the same simulation parameters that were used to generate in Fig. 1 panel (a1). Predictably, in the slow kinetics regime, we expect both HMJP and HMM to do well and this is what we find.

In Fig. A.1 panel (a3), we observe that there is at most one switch during one integration period. We see in Fig. A.1 panel (a4) that both HMJP and HMM MAP trajectory estimates are capable of capturing these switches in the true trajectory.

Next, we move to the posterior probability distribution over state levels shown in Fig. A.2 panels (a1)-(a2). In panel (a1), we observe that the HMM grossly overestimates (by about 30%) *μ*_*σ*_1__ and in panel (a2), HMM underestimates (about 3%) *μ*_*σ*_2__. By contrast, the HMJP posterior probability distribution for the state levels provide very good state level estimates for both *μ*_*σ*_1__ and *μ*_*σ*_2__. Put differently, true *μ*_*σ*_1__, *μ*_*σ*_2__ values occur in the 95% confidence intervals of the HMJP posterior distributions over state levels.

Finally, we discuss the posterior distribution over transition probabilities obtained from HMJP and HMM frameworks in Fig. A.4 panels (b1)-(b4). We expect both HMM and HMJP to provide very good estimates for probabilities of switching among the states as these are associated with slow switching rates prescribed with *λ*_*σ*_1_→*σ*_2__, *λ*_*σ*_2_→*σ*_1__ (see Eq. (31)).

In Fig. A.2 panels (b1)-(b4), we observe that HMM and HMJP perform quite well in estimating transition probabilities based on its narrow 95% confidence intervals.

Now we move on to the results for mixed switching rates. We define the values for the parameters *τ_f_*, *τ* as we proceed.

#### A.1.2 Comparison of HMJP with HMM for Mixed Switching Rates

In this section, we test the performance of HMJPs and HMMs in estimating trajectory Fig. A.3, state levels Fig. A.4 and transition probabilities Fig. A.4 on the analysis of the simulated measurements associated with the switching rates *λ*_*σ*_1_→*σ*_2__, *λ*_*σ*_2_→*σ*_1__, with a parameter *τ_f_* which sets the timescale of the system kinetics,

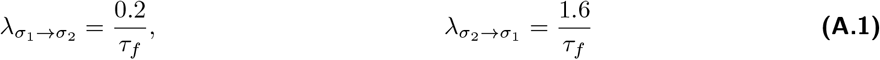

**Fig. A.1.**
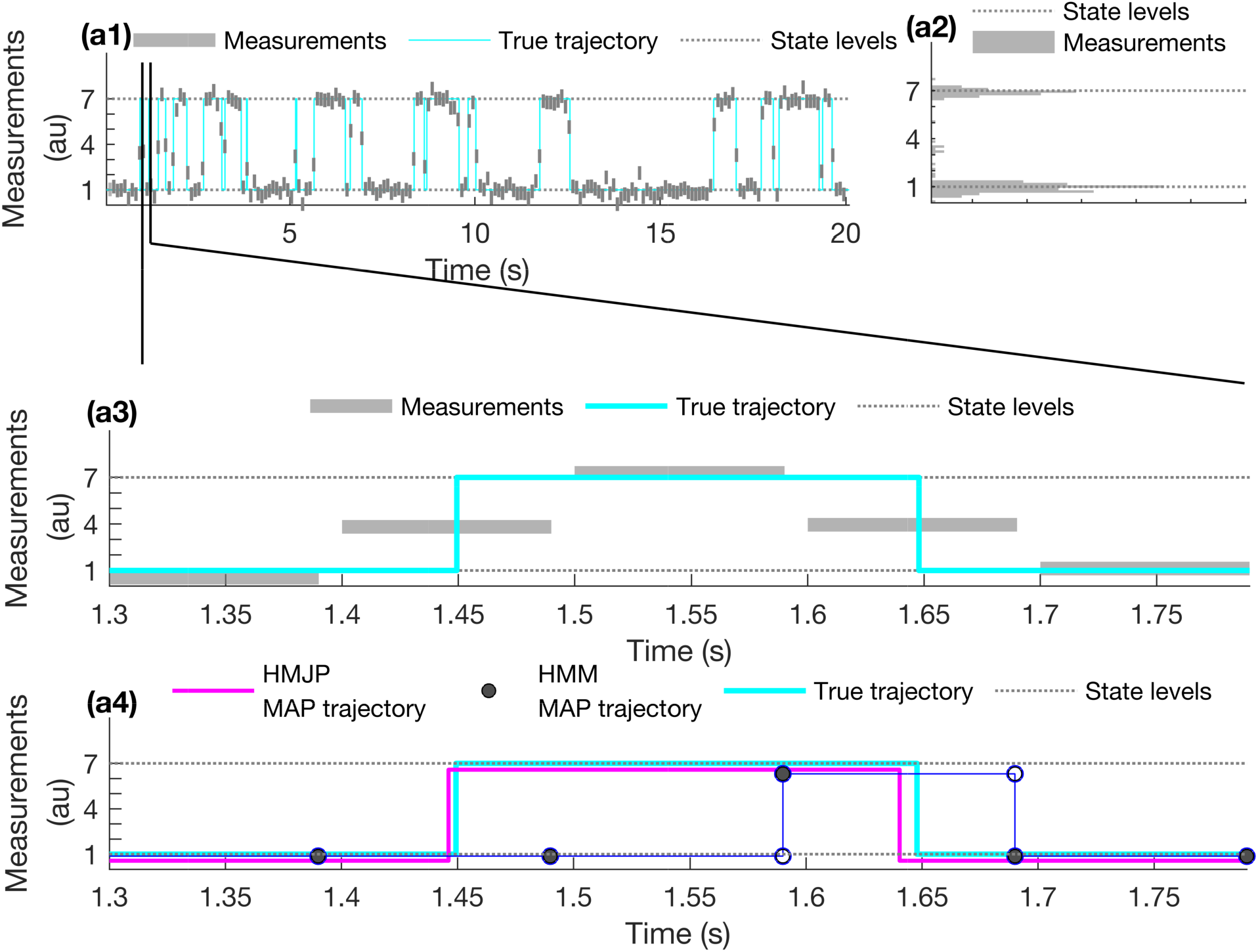
HMJP with HMM trajectory estimates for slow switching. Here we provide trajectory estimates obtained with HMJPs and HMMs. The results for the HMJP were previously provided in Figs. 2 and 3 but here we add the results for the HMM in panel (a4). We follow identical coloring schemes and convention as with Fig. 2 of the main. The parameters used are identical to those of Fig. 2.

**Fig. A.2.**
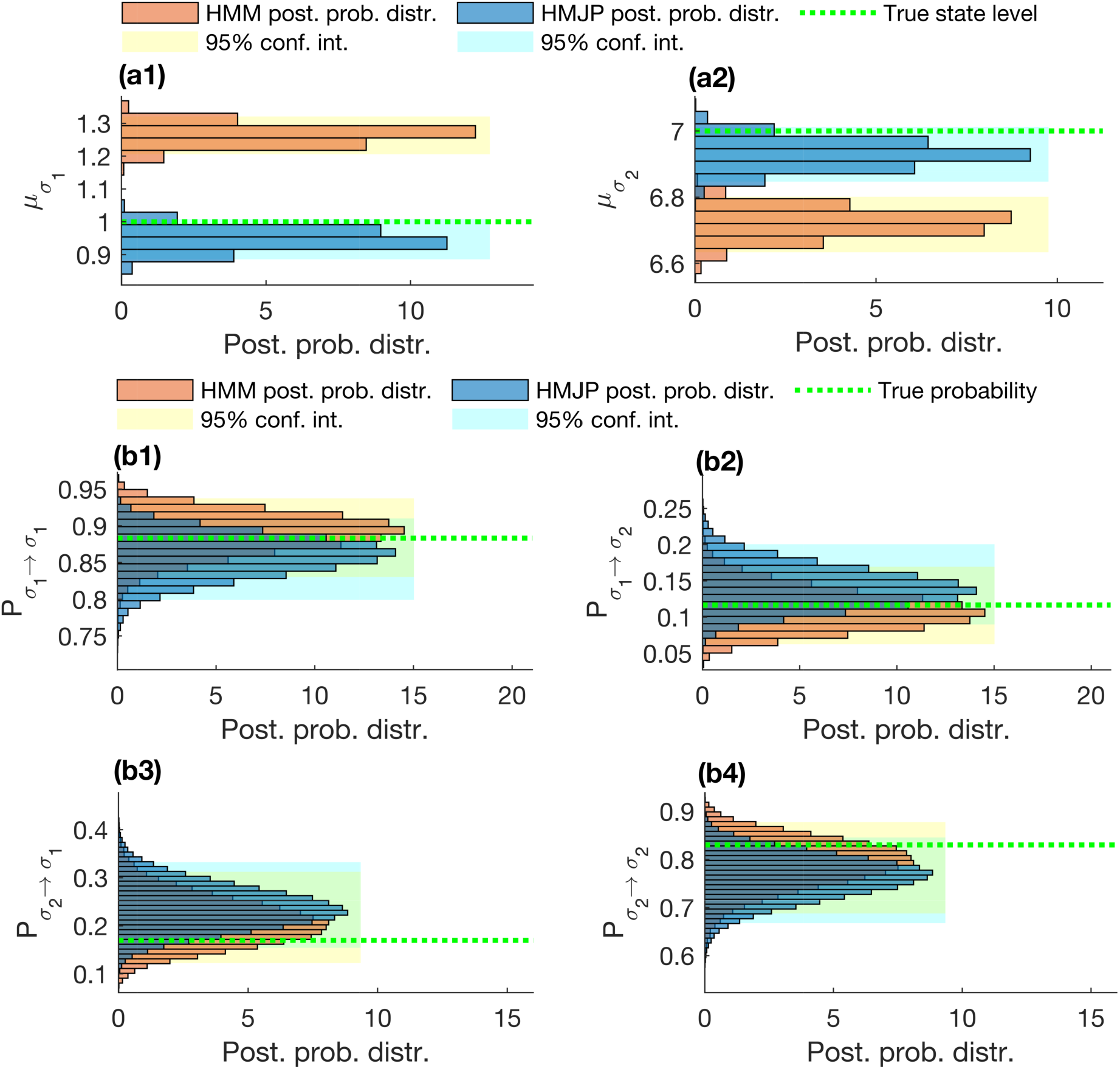
HMJP with HMM state level and transition probability estimates for slow switching. Here we provide posterior state level and transition probability estimates obtained with HMJPs and HMMs. The results for the HMJP on posterior state levels were previously provided in Fig. 3 but here we add the results for the posterior transition probabilities for both HMJP and HMM in panels (b1)-(b4) and posterior state level estimaets for HMM in panels (a1)-(a2). We follow identical coloring schemes and convention as with Fig. 3 of the main. The parameters used are identical to those of Fig. 2.

**Fig. A.3.**
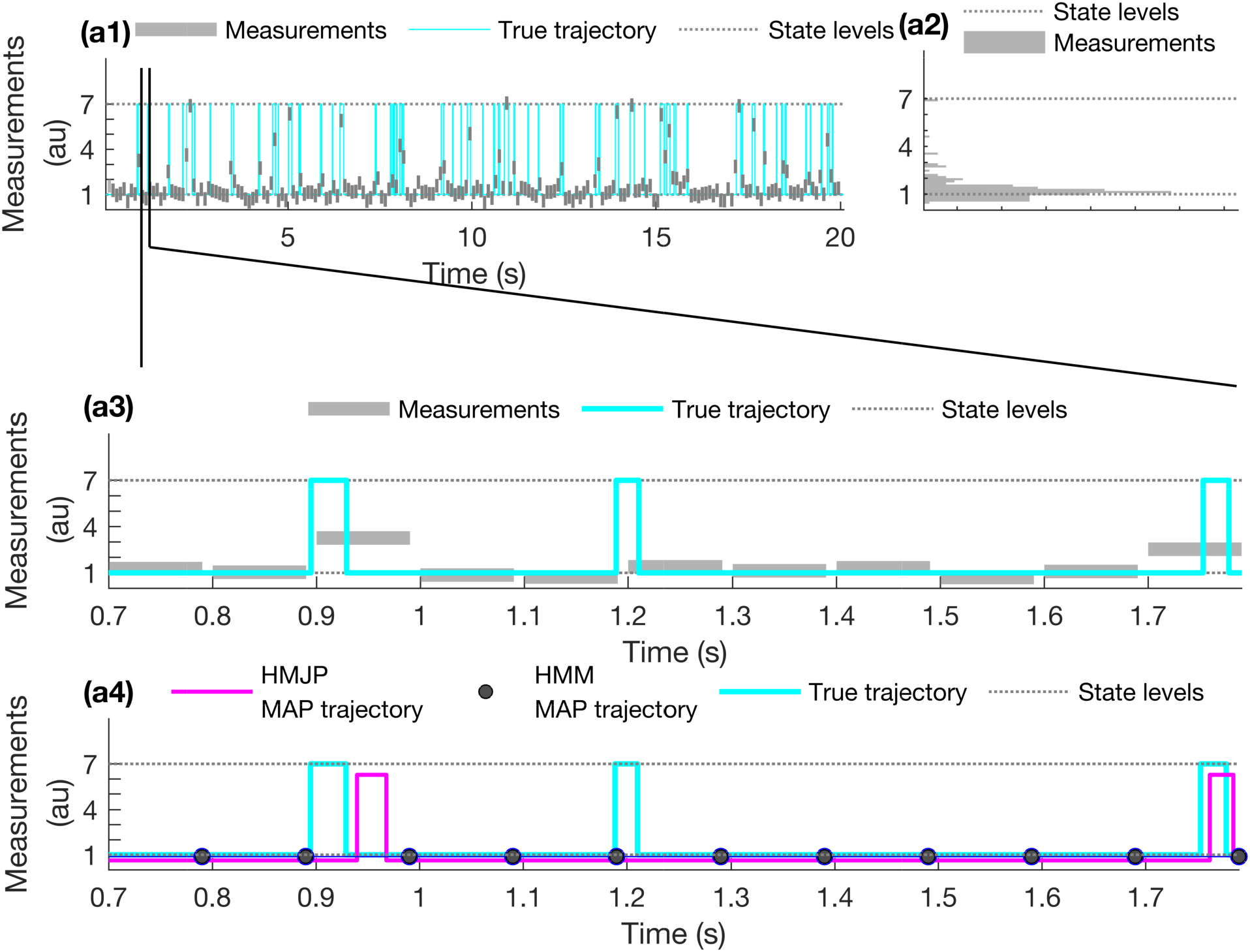
HMJP with HMM trajectory estimates for mixed switching. Here we provide trajectory estimates obtained with the HMJP and HMM when switching rates are slower or faster than the data acquisition rate, 1/Δt = 10 (1/s). We follow identical coloring schemes and convention as in Fig. A.1. We see that the HMJP MAP trajectory is able to capture most of the fast switches between the states σ_1_ and σ_2_. Simulated measurements are generated with λ_σ_1_→σ_2__, λ_σ_2_→σ_1__ (see Eq. (A.1)) where the data acquisition happens every Δt = 0.1 s with τ_f_ = 1/15 s and τ = 0.09 s starting at t_0_ = 0.05 s until t_N_ = 20 s.

where *τ_f_* = 1/15 s and data acquisition rate, 10 (1/s). We start with Fig. A.3 panel (a3) where we observe that there are multiple fast switching events during one integration period. We see in Fig. A.3 panel (a4) that the HMJP MAP trajectory estimate is capable of capturing most of the fast switches occurring in the true trajectory while the HMM MAP trajectory estimate provides a poor trajectory estimate as compared to ground truth.

The HMM’s poor MAP trajectory estimate can be explained with the posterior probability distribution over state levels shown in Fig. A.4 panels (a1)-(a2). In Fig. A.4, we provide the superposed posterior distributions for HMJP and HMM over state levels in panels (a1)-(a2) and transition probabilities in panels (b1)-(b4). In panel (a1), we observe that the HMM grossly overestimates (by about 30%) *μ*_*σ*_1__. In panel (a2), the HMM grossly underestimates (by about 20%) *μ*_*σ*_2__. By contrast, the HMJP posterior probability distribution for the state levels provide very good state level estimates for both *μ*_*σ*_1__ and *μ*_*σ*_2__. Put differently, true *μ*_*σ*_1__, *μ*_*σ*_2__ values take place at the 95% confidence intervals of the HMJP posterior distributions over state levels. Finally, we discuss the posterior distribution over transition probabilities obtained from the HMJP and HMM frameworks Fig. A.4 panels (b1)-(b4). We expect to see HMMs provide very good estimates for the transition probabilities from state *σ*_1_ to itself and *σ*_2_ as these are associated with slow switching rates prescribed with *λ*_*σ*_1_→*σ*_2__ (see Eq. (A.1)). By contrast, we expect wide posterior distributions over transition probabilities from conformational state *σ*_2_ to itself and *σ*i as *P*_*σ*_2_→*σ*_1__ and *P*_*σ*_2_→*σ*_2__ are associated to the fast switching rates defined by *λ*_*σ*_2_→*σ*_1__ (see Eq. (A.1)). In Fig. A.4 panels (b1)-(b4), we observe that the HMJP still performs quite well in estimating transition probabilities based on its narrow 95% confidence intervals. Next, we provide the posterior distributions over rates obtained by the HMJP for fast and mixed switching kinetics.

#### A.1.3 Robustness Analysis for HMJP Posterior Rate Estimates for Fast and Mixed Kinetics

In the main text, for sake of comparison between HMJPs and HMMs, we compared their posterior estimates for the transition probability. However, HMJPs are also capable of providing estimates for the transition rates, not just transition kinetics.

Using parameter values to simulate the results of Fig. 4 for the fast kinetics and Fig. A.3 for the mixed rates we show rate posteriors below in Fig. A.5.

In Fig. A.5, we observe that the HMJP is capable of providing very good estimates for the analyzed data generated with slow, fast and mixed switching rates. For all simulations, we always have the same shape (*η*) and scale 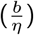 hyperparameters for the gamma prior distributions and we set *η* = 2 and *b* = 300. Now we move on to the rest of the additional results. We define the values for the parameters *τ_f_, τ* as we proceed.

#### A.1.4 Robustness Analysis with Respect to Data Set Length for HMM Posterior State Level and Transition Probability Estimates

Here we show how our posterior estimates over quantities including MAP trajectory estimate, state level and transition probabilities vary as a function of variable data set length, see Figs. A.6 and A.7.

We focused on the case of fast kinetics because we want to demonstrate that the poor estimates are not due to finiteness of data. The parameters used are identical to those used to generate the data for Fig. 1 panel (b1) except for the final time that is *t_N_* = 60 s.

We provide a quantitative metric (area under the curve (102); see Eq. (A.2)) for deviation from ground truth in Fig. A.9.

**Fig. A.4.**
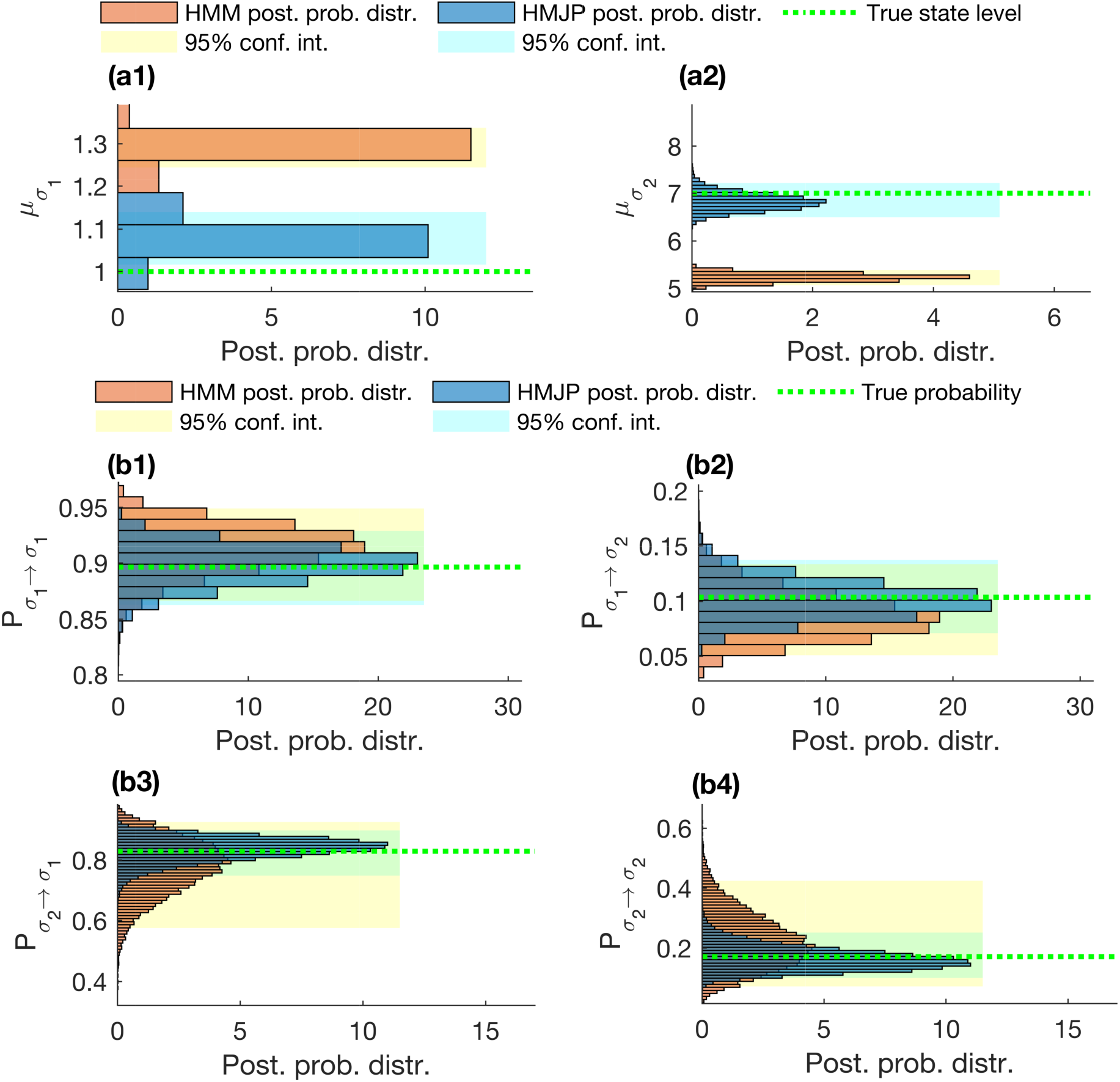
HMJP with HMM state level and transition probability estimates for mixed switching. Here we provide posterior state level and transition probability estimates obtained with the HMJP and HMM when the switching rate is slower or faster than the data acquisition rate, The results for the HMJP and HMM MAP trajectory estimates are provided in Fig. A.3. We show the superposed posterior state level estimates for the HMJP and HMM in panels (a1)-(a2) and the superposed results for the posterior transition probabilities for both HMJP and HMM in panels (b1)-(b4). We follow identical coloring schemes and convention as with Fig. 3 of the main text. Here, simulated measurements are generated with the same parameters as those provided in Fig. A.3.

**Fig. A.5.**
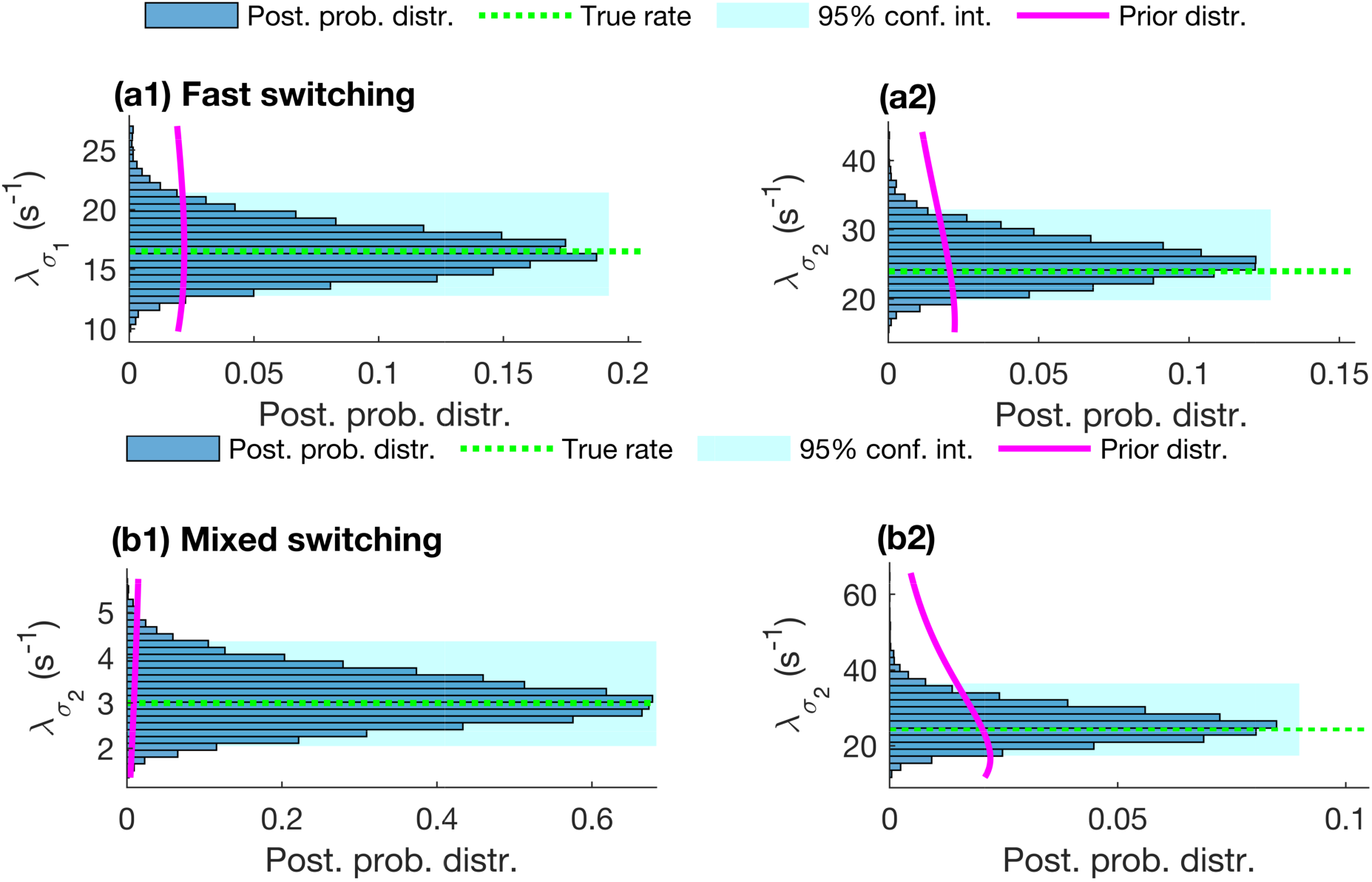
HMJP rate estimates for fast and mixed switching rates. Here we provide posterior rate estimates with the HMJP for the simulated measurements provided in Figs. 4 and A.3. We follow identical coloring schemes and convention as with Fig. 3 of the main. Here, simulated measurements associated with estimates given in panels (a1)-(a2) and (b1)-(b2) are generated with the same parameters used in Figs. 2 and 4.

**Fig. A.6.**
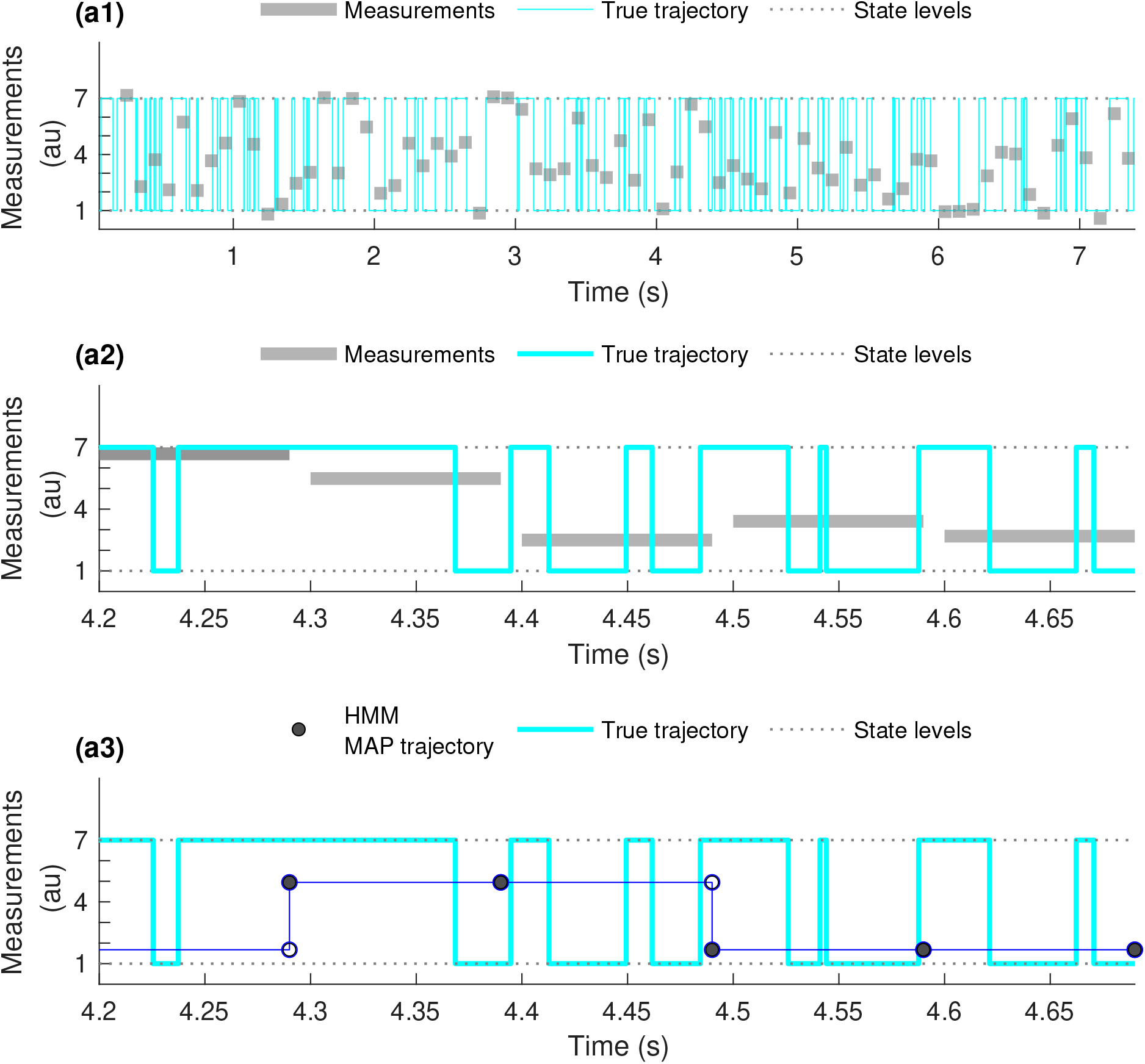
HMM trajectory estimates for fast state switching. Here we provide trajectory estimates obtained with the HMM when the switching rates are faster than the data acquisition rate, 1/Δt = 10 (1/s). We follow identical coloring schemes and conventions as we do in Fig. A.1. We demonstrate that the HMM MAP trajectory poorly estimates the true trajectory.

**Fig. A.7.**
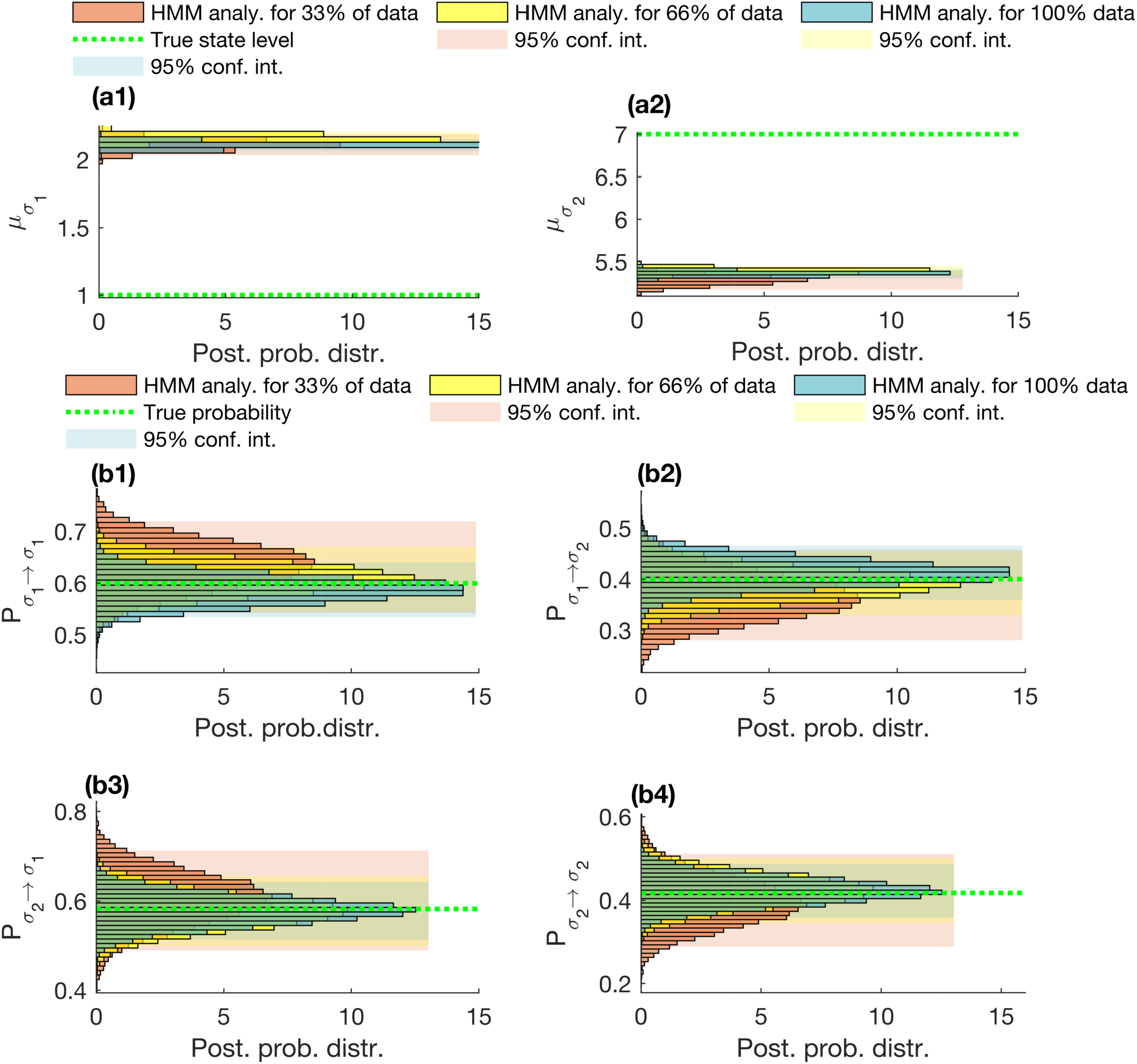
Finiteness of data analysis for HMM state level and transition probability estimates in the presence of fast state switching. Here, we would like to show the effect of set data set length on the state level and transition probability estimates provided by HMMs on the simulated measurements shown in Fig. A.6. To begin, in all panels, we see the superposed posterior distributions of the state levels and transition probabilities obtained from HMMs for 33% (orange), 66% (yellow), 100% (blue) of the entire simulated measurements along with their associated 95% confidence intervals, the true state levels and transition probabilities (green dashed line).

We performed our investigation for the HMM in 3 parts. First, we analyzed 33% of the data, next 66% of the data and lastly we analyzed the entire data set with the HMM. We showed the simulated trajectory for the first 12% of the entire data set in Fig. A.6. In Fig. A.6 panel (a1), we show the true trajectory (cyan) and the measurements (gray rectangles). Next, in Fig. A.6 panel (a2) we showed the zoomed true trajectory and measurements. Finally, in panel (a3), we provide the superposed true trajectory with HMM MAP trajectory estimate. We observe that the HMM MAP trajectory estimate is a poor trajectory estimate. This poor trajectory estimates of HMM can be explained by the poorly estimated state levels, see Fig. A.7 panels (a1)-(a2). In Fig. A.7, we observe in both panels (a1)-(a2) that more data does not provide better state level estimates for HMMs. Subsequently, in Fig. A.7 panels (b1)-(b4), we demonstrate that the posterior distributions over transition probabilities obtained with HMM for all 3 data sets do not differ greatly from each other. Next we investigate the HMJP’s performance in estimating transition probabilities with and without pre-specifying the ground truth trajectory.

#### A.1.5 Robustness Analysis with Respect to Learning the Trajectory Simultaneously with Transition Probability Estimates for HMJPs

In this section, we test the effect of estimating trajectories simultaneously with transition probability estimates using HMJPs. With sufficient data, we expect HMJPs to perform similarly well when: 1) the trajectory is assumed to be known; and 2) the trajectory is not assumed to be known and thus to be estimated as well. We demonstrated that posterior transition probabilities look similar when the trajectory is assumed to be known or not known in Fig. A.8.

Now we compare the performance of HMJP and HMM in estimating trajectories based on the average area under their posterior trajectory estimates.

#### A.1.6 Comparison of the Posterior Trajectory Estimates of HMM and HMJP

In this section, we provide a metric to compare the quality of the trajectory determination by HMMs and HMJPs (102). As we have a posterior distribution over trajectories 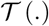 obtained by HMMs and HMJPs, we also have posterior distributions over areas

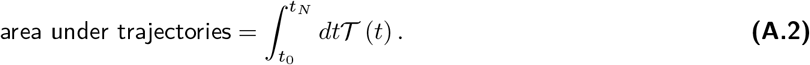

In Fig. A.9, we compare the mean point statistic of *area under trajectories* for both HMM and HMJP.

We carried out this analysis for the simulated measurements investigated in the main text generated with slow switching rates Fig. 2 and fast switching rates Fig. 4.

We observe that, the average areas under HMJP posterior trajectory estimates are very closed to the areas under the true trajectories for slow and fast switching rate cases.

**Fig. A.8.**
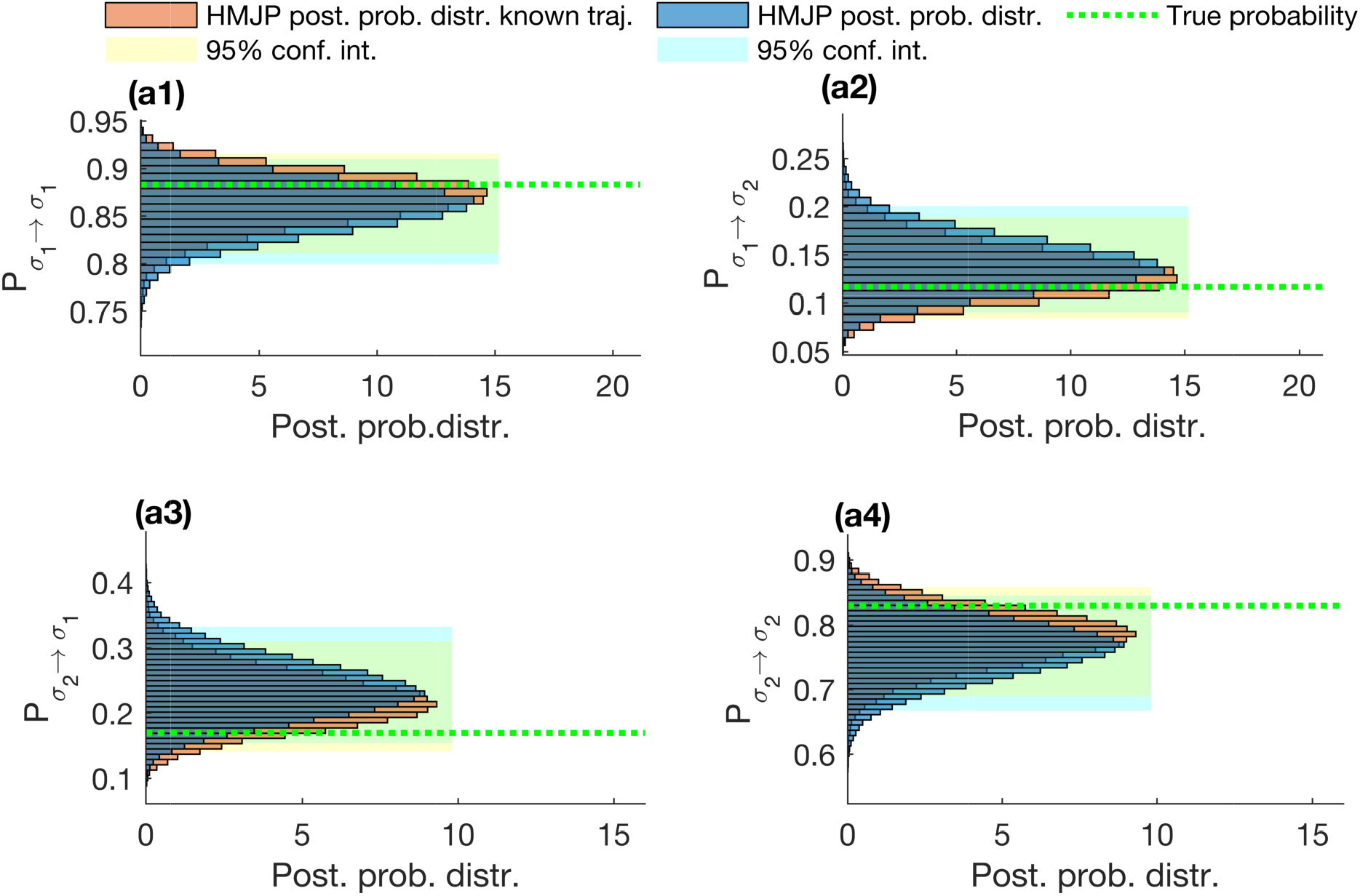
HMJP transition probability estimates with and without trajectory information. Here, we provide the analysis of the simulated measurements provided in Fig. 2 panel (a1). In this figure, we illustrate the HMJP’s performance in estimating transition probabilities with and without learning the trajectory. To do so, in each figure panel we provide the superposed posterior distributions over transition probabilities labeled as P_σ_k_→σ_k′__ for all k,k′ = 1, 2 obtained by HMJP with unknown system trajectory (blue), with known system trajectory (orange) along with their associated 95% confidence intervals for the estimates and the true transition probabilities (dashed green lines). We observe that there is very limited change in the posterior transition probabilities based on their 95% confidence intervals when the trajectory is known or, alternatively, learned simultaneously.

**Fig. A.9.**
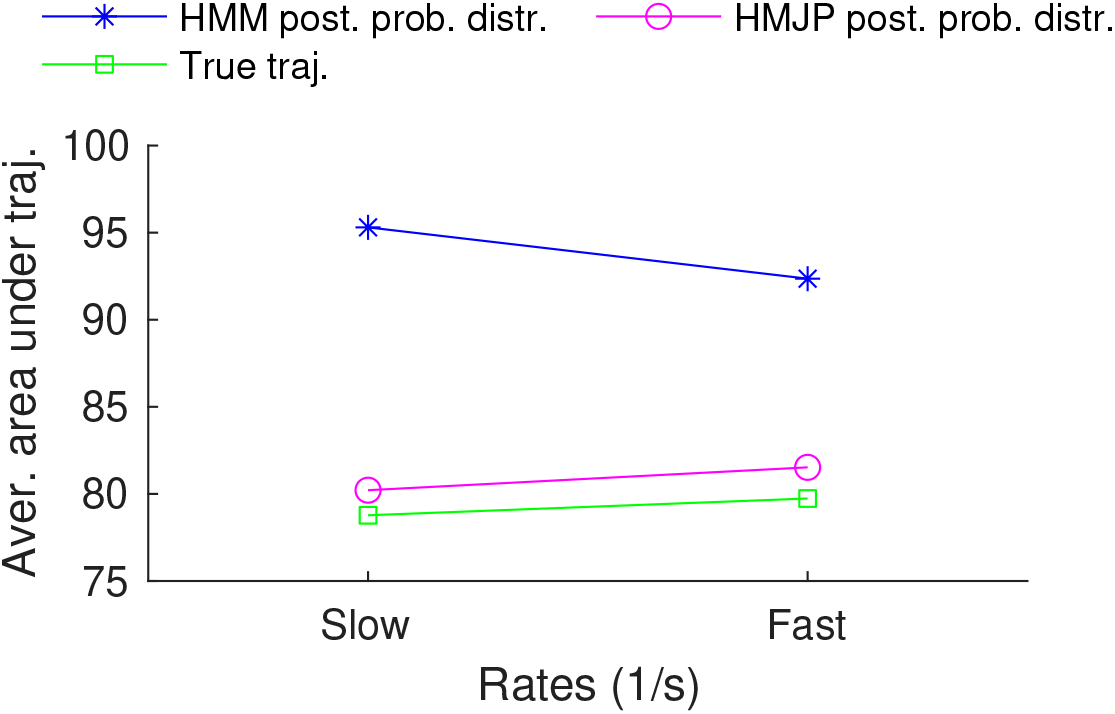
HMJP with HMM average area under the trajectory estimates. Here we compare the performance of HMJPs and HMMs in estimating trajectories based on the average area under their posterior trajectory estimate for the simulated measurements generated with slow switching rates (denoted with “Slow”), see Fig. A.1 and fast switching rates (denoted with “Fast”), see Fig. 4. In this figure, we provide the average areas under the HMM (blue star) and HMJP (magenta circle) posterior trajectory estimates and the area under the true trajectory (green circle) for both simulated measurements generated with slow switching rates and fast switching rates.

### A.2 Detailed Description of the Statistical Models

#### A.2.1 Hidden Markov Jump Process

##### A.2.1.1 Summary of Equations

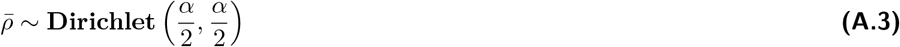

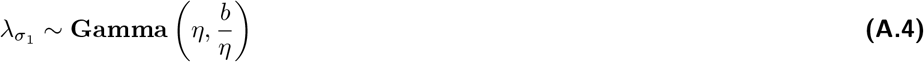

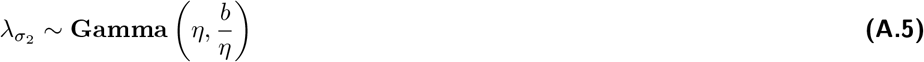

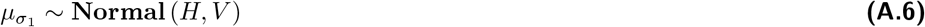

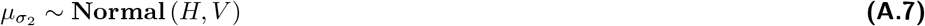

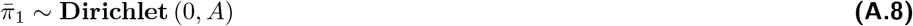

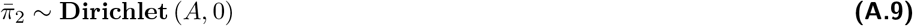

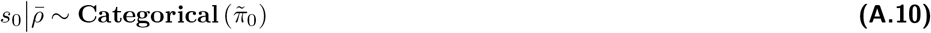

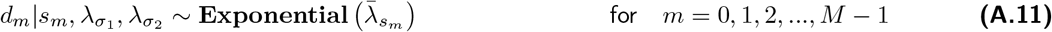

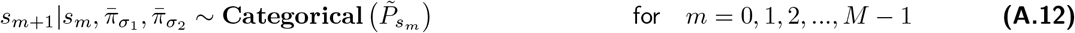

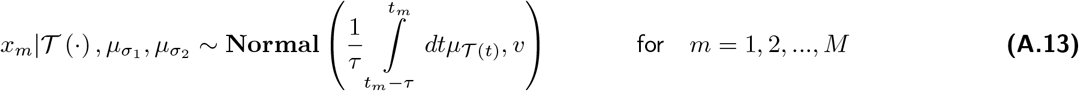

where 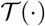 is formulated as follows

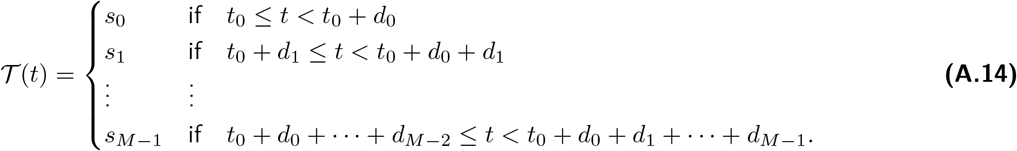

with *M* determined based on the first time

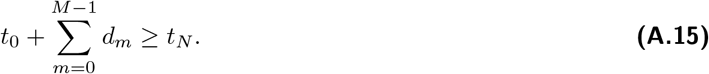

In this manuscript as in the case of HMM framework, we set *α* = *A* = 1, FWHM = 0.25, *H* = mean(data) au, and *V* = 1 au.

##### A.2.1.2 Description of the Computational Scheme

Our MCMC exploits Gibbs sampling scheme (38, 39, 81). Accordingly posterior samples are generated by updating each variables involved sequentially by sampling conditioned on all other variables and the measurements **x**. Conceptually, the steps involved in the generation of each posterior 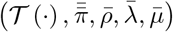 involve

1. Updating the trajectory 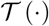,
2. Updating the transition probability matrix 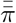 and 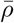,
3. Updating 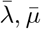.

Specifically, for step 1, to sample the trajectory 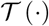, we need two different strategies. As the state dynamics of the HMM evolve in discrete time, we use a *forward filtering backward sampling* scheme (34, 38–43). For the HMJP, as the state dynamics evolve in continuous time, we perform *uniformization* prior to sampling a new trajectory (62, 69, 70, 103, 104) and then we perform Gibbs sampling to obtain *s*_1:M_.

For steps 2, 3 we have conjugate priors. Thus, we can do direct sampling.

It is in step 1, where we update the trajectory of the system, where the HMM and HMJP differ most in methodology and computational cost. This is because of the *uniformization* (103) required of the HMJP that sets *M* as described in Eq. (8).

#### A.2.2 Hidden Markov Model

##### A.2.2.1 Summary of Equations

For *K* = 2, the full set of HMM equations is

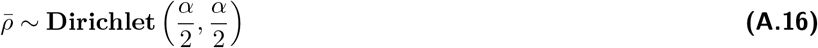

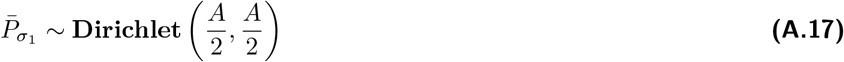

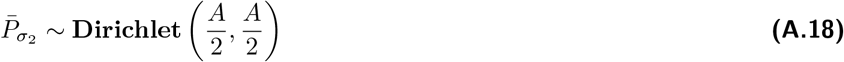

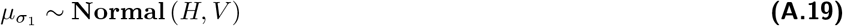

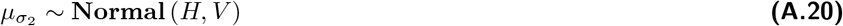

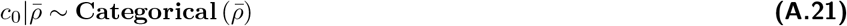

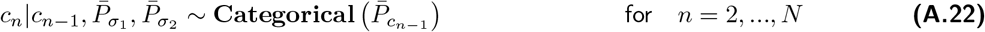

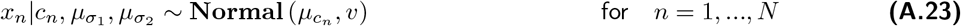

##### A.2.2.2 Description of the Computational Scheme

The joint probability distribution of our framework is 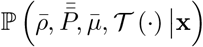. As we do not have conjugacy between dynamics and the measurements, it is not possible to have a direct sampling from the posterior distribution. For this reason, we develop a specialized Markov Chain Monte Carlo (MCMC) scheme that can be used to generate pseudorandom samples (81). This scheme is explained in detail below. A working implementation of the resulting scheme in source code and GUI forms provided through the author’s website.

Our MCMC exploits Gibbs sampling scheme (38–41, 43, 81). Accordingly posterior samples are generated by updating each one of the variables involved sequentially by sampling conditioned on all other variables and the measurements **x**. Conceptually, the steps involved in the generation of each posterior 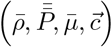 are

1. Update the trajectory 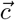,
2. Update transition probabilities 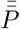,
3. Update transition probabilities 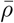,
4. Update transition probabilities 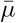.

Sampling the trajectory 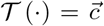 involves forward filtering-backward sampling scheme (38, 39). Updating the transition probability matrix can be carried out for every row separately on account of the independence of the Dirichlet prior on each 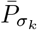 and 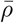. Because of the conjugacy between the Dirichlet distribution and Categorical distribution, we have direct sampling updates for all 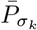 and 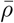 (81). Similarly, due to conjugacy of the Normal distribution with itself, we also have direct updates for *μ_σ_k__* for all *k* = 1, 2.

We mention that updating *c* is carried out via forward filtering backward sampling (81) approach (38, 39, 43, 81).

In the process of generating pseudorandom numbers from the posterior distribution described above, the first ~ 1000 pseudorandom numbers are discarded to account for MCMC burn-in. The rest of the generated numbers contribute to the posterior probability distribution over the trajectories, transition probability matrix and state levels.

Next, we provide the detailed information regarding these listed updates mentioned above.

##### A.2.2.3 Overview of the Sampling Updates

###### 1. Sampling a trajectory 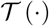

We will sample continuous time trajectories using *uniformization* (62, 69, 103) whose key steps 1, 2, 3 are illustrated in Fig. A.10. Uniformization uses ideas from discrete time-discrete state space Markov processes.

To sample new trajectories, we start from rates, see Eq. (A.1) and old trajectories (see Fig. A.10 panel (a)) generating these trajectories using a Markov jump process. The trajectories are characterized by states 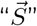 and associated holding times,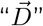 namely,

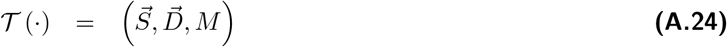

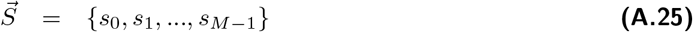

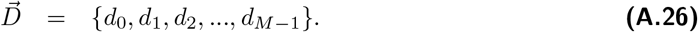

Following Eq. (13) of the main, we can construct a generator matrix 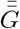. As a reminder, the system occupies each state *s_m_* during the interval [*t_m_, t*_*m*+1_) with *d_m_* = (*t*_*m*+1_ – *t_m_*) holding times for all *m* = 0,1, 2,…,*M* – 1. As we can see from the cartoon figure in Fig. A.10 panel (b), we must now add virtual jumps. At the *m^th^* time interval, we add a number virtual jumps and distribute these *uniformly* in *t*_*m*+1_ – *t_m_*. We select the number of jumps according to a Poisson distribution with intensity 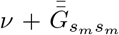. Here 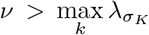. To achieve this, we set 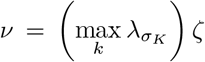. In this manuscript, we choose *ζ* = 2 as suggested in (62).

As such we initially overestimate the number of jump locations. We subsequently need to prune these down by determining if these virtual jumps coincide with real jumps. In principle we can set v to as large a value as we like, this only increases the computational cost of pruning.

**Figure A.10.**
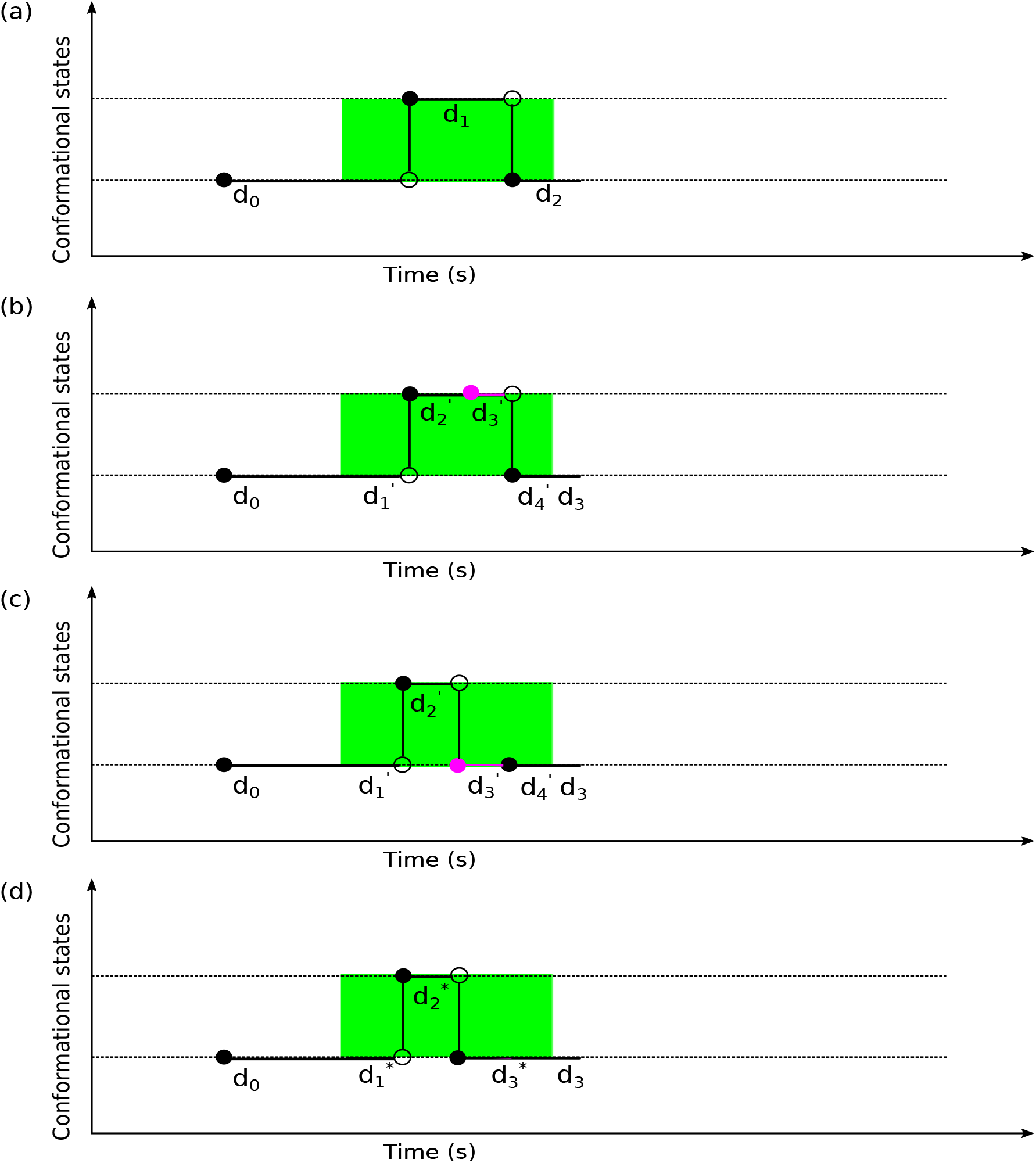
A conceptual description of uniformization on sampling a Markov jump process. Here, we explain how we perform uniformization to obtain a new trajectory estimate using the HMJP in 4 panels starting from an old trajectory estimate. In panel (a), we show the old trajectory estimate that we would like to update. This green region is associated with the n^th^ integration period that give rise to measurement x_n_. We note that there is 1 jump in this old trajectory estimate with 2 holding times that are labeled with d_1_, d_2_. We simply define the states associated with d_0_, d_1_, d_2_, d_3_ as s_0_, s_1_, s_2_, s_3_, respectively. In panel (b), we introduce virtual jump events with circles in magenta and then we relabel the trajectory with new holding times and states. Upon relabeling, we now have states 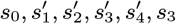 associated with the holding times 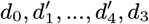. Next, we apply Gibbs sampling to sample each 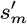 for all m = 1, 2, 3, 4 conditioned on all other variables. We show the newly sampled states in panel (c). The last step in this uniformization is to remove the self-transitions from the new sampled trajectory so that it remain a Markov jump process. To do so, we remove the self-transitions and in panel (d), we show the new sampled trajectory that is a Markov jump process with states 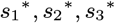 and associated holding times 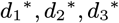, respectively.

To prune, as shown in panel (b) of Fig. A.10, we now must revert to a discrete Markov chain picture. We begin by defining the transition matrix 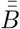 that coincides with the Poisson intensity of 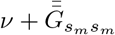. It is

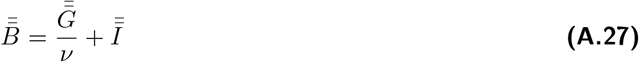

Given a transition matrix 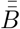, we can now straightforwardly apply the forward filter-backward sampler to sample those states visited to sample the states at each virtual jump point.

Below we provide the details of sampling a part of a trajectory that is associated with observation *x_n_* with Gibbs sampling scheme.

First, we write down the target distribution and then we factorize it as follows

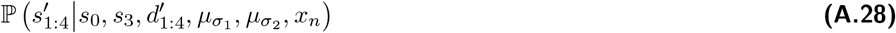

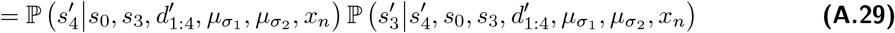

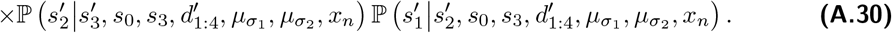

Next, we have to sample each part of this target distribution given above according to the following

a. 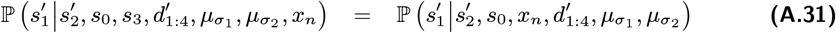

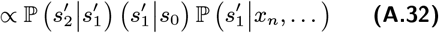

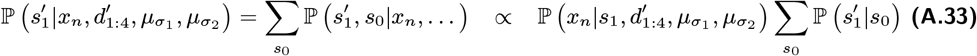
b. 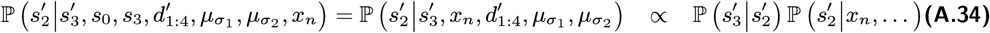

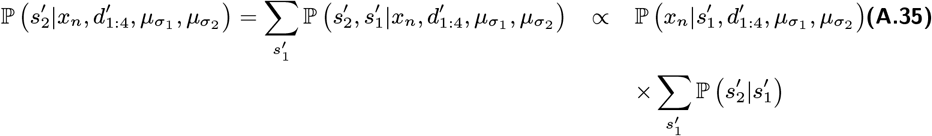
c. 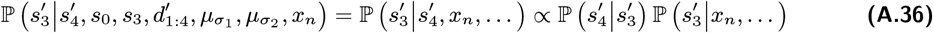

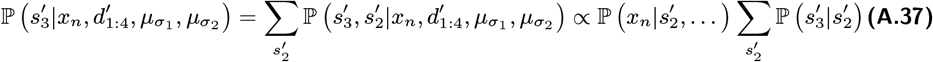
d. 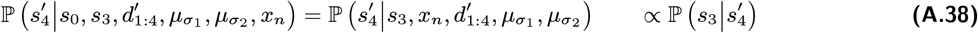

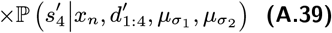

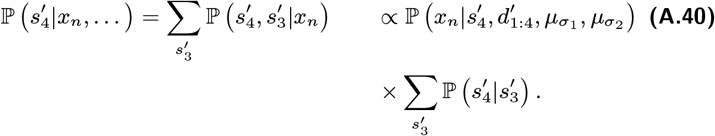

We repeat this process of sampling trajectory for all observations *x_n_* where *n* = 1, 2,…, *N*. We should note that 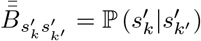 with *k, k*’ = 1, 2, 3, 4.

Once we obtain the new trajectory (see Fig. A.10 panel (c)) we observe that this new trajectory includes self transition events at the jump times.

However, occupying the same steps after the jump times is not allowed for continuous time processes, therefore we can not have self transition events in Markov jump processes. To obtain a Markov jump process from this sampled new trajectory, we drop the self transition events (see Fig. A.10 panel (d)) thereby obtaining a new trajectory which is a Markov jump process.

Next we explain how to update the transition probability matrix 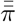.

###### 2. Sampling transition probabilities 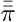 and 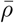

We placed conjugate Dirichlet distributions to the Categorical distribution for the rows of 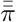. Therefore, we update each row of 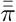 via direct sampling as we do for the case of the HMM

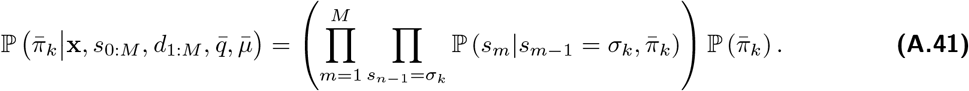

Similar direct sampling formulation applies to 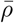 as follows

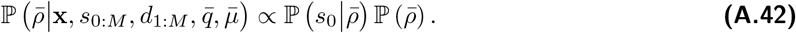

Next, we provide the details of sampling 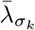 for all *k* =1, 2.

###### 3. Sampling 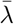

We placed conjugate Gamma prior distributions to the Exponential distribution for 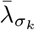 where *k* =1, 2. Therefore updating all 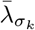 with *k* =1, 2 is carried out with direct sampling based on the following formulation

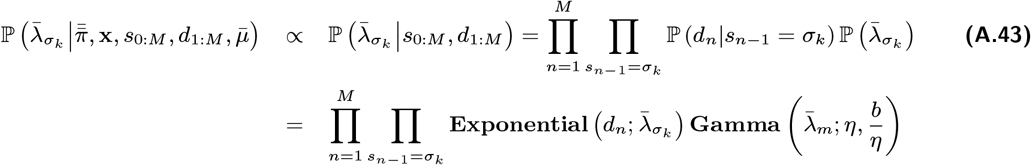

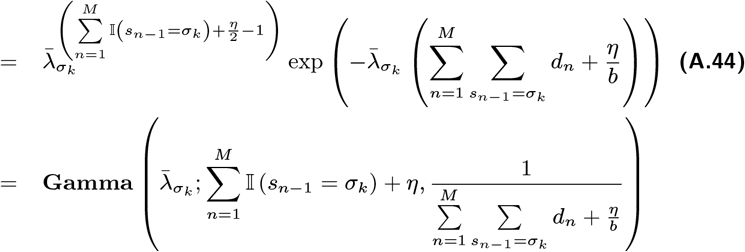

for *k* = 1, 2. In this formulation, 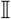 denotes the indicator function and for example 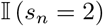 is 1 if *s_n_* = 2 for some *n* = 1, 2,…, *M* and 0 otherwise.

###### 4. Sampling 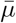

We perform the updates of *μ_σ_k__* for all *k* = 1, 2 as a part of the Gibbs sampling scheme. Therefore, we need the full conditional distributions of *μ_σ_k__* for all *k* = 1, 2 these are 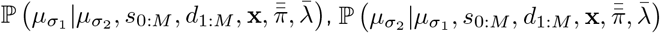. Below, we provide the explicit update formula for *μ*_*σ*_1__ and the similar for-mulation holds for *μ*_*σ*_2__.

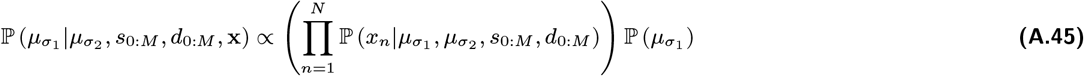

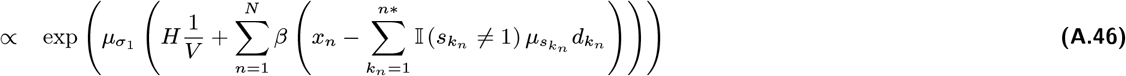

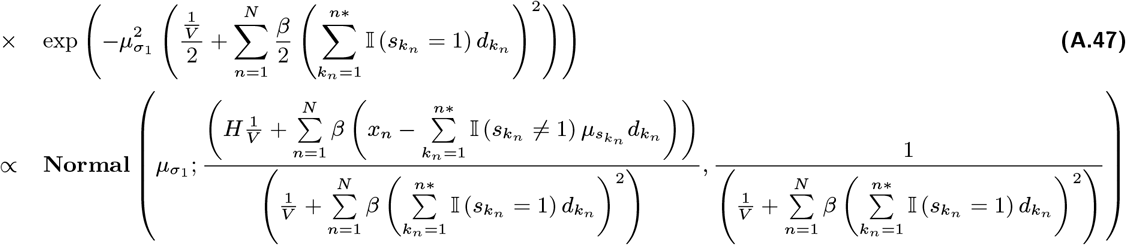

where **x** = (*x*_1:*N*_).

### A.3 Notation

**Table A.1.**
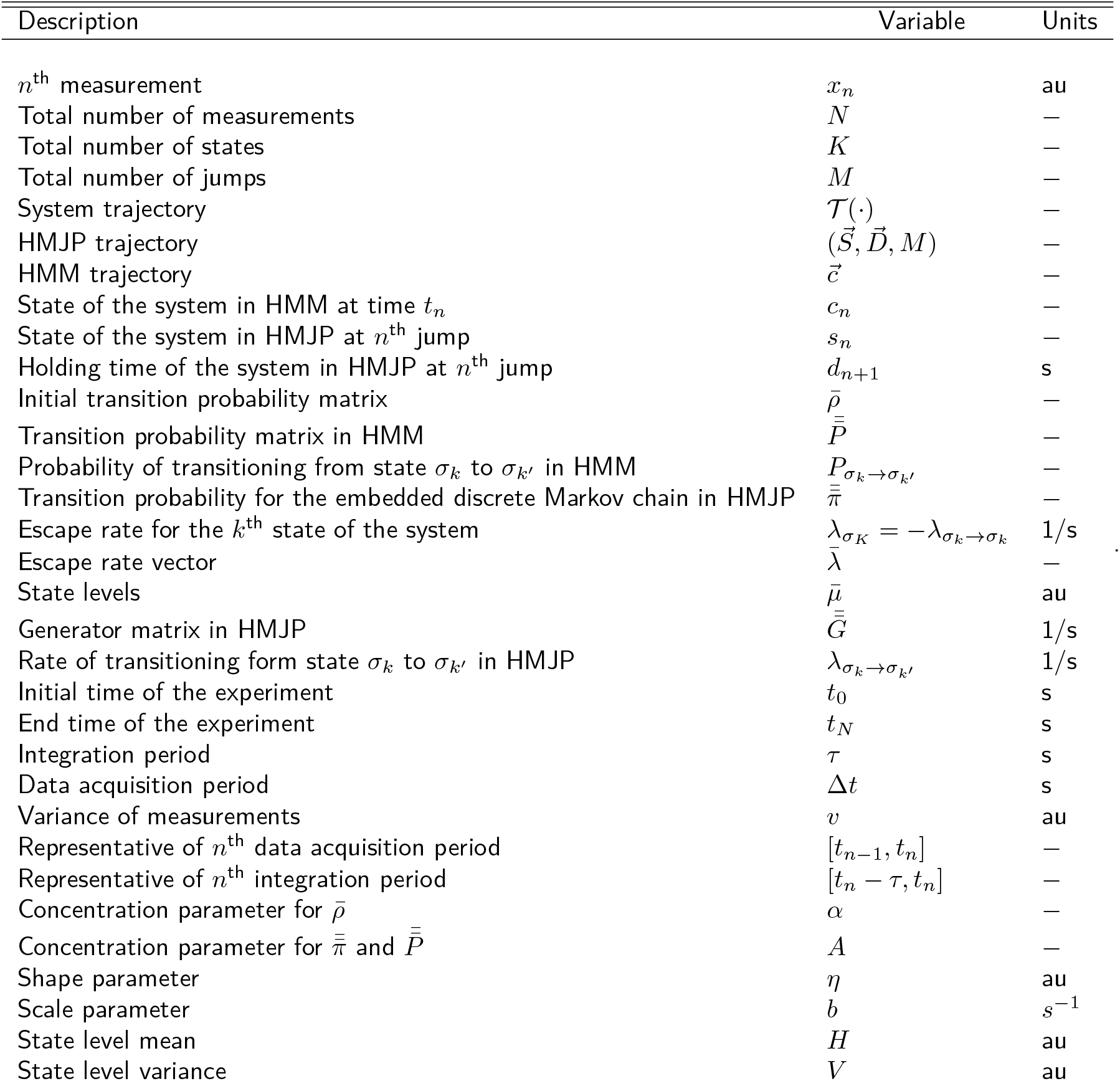
Notation conventions

**Table A.2.** Parameter choices and units

| Variable | $H$ | $V$ | $\eta$ | $b$ | $A$ | $\alpha$ |
| --- | --- | --- | --- | --- | --- | --- |
| Units | au | au | au | $s^{-1}$ | - | - |
| Fig. 2 | mean of measurements | 1 | 2 | 300 | 1 | 1 |
| Fig. 3 | mean of measurements | 1 | 2 | 300 | 1 | 1 |
| Fig. 4 | mean of measurements | 1 | 2 | 300 | 1 | 1 |
| Fig. 5 | mean of measurements | 1 | 2 | 300 | 1 | 1 |
| Fig. A.1 | mean of measurements | 1 | 2 | 300 | 1 | 1 |
| Fig. A.2 | mean of measurements | 1 | 2 | 300 | 1 | 1 |
| Fig. A.3 | mean of measurements | 1 | 2 | 300 | 1 | 1 |
| Fig. A.4 | mean of measurements | 1 | 2 | 300 | 1 | 1 |
| Fig. A.6 | mean of measurements | 1 | 2 | 300 | 1 | 1 |
| Fig. A.7 | mean of measurements | 1 | 2 | 300 | 1 | 1 |
| Fig. A.8 | mean of measurements | 1 | 2 | 300 | 1 | 1 |
| Fig. A.5 | mean of measurements | 1 | 2 | 300 | 1 | 1 |

